# Artificial upwelling leads to a large increase in surface dissolved organic matter concentrations

**DOI:** 10.1101/2022.06.27.496799

**Authors:** Markel Gómez-Letona, Marta Sebastián, Isabel Baños, María Fernanda Montero, Clàudia Pérez Barrancos, Moritz Baumann, Ulf Riebesell, Javier Arístegui

## Abstract

In the face of climate change there is a need to reduce atmospheric CO_2_ concentrations. Artificial upwelling of nutrient-rich deep waters has been proposed as a method to enhance the biological carbon pump in oligotrophic oceanic regions in order to increase carbon sequestration. However, the fate of the newly produced organic matter, and specifically of its resulting dissolved fraction, is not clearly understood. Here we examine the effect of different upwelling intensities and modes (single pulse versus recurring pulses) on the dissolved organic matter pool (DOM). We introduced nutrient-rich deep water to large scale mesocosms (∼44 m^3^) in the oligotrophic subtropical North Atlantic and found that artificial upwelling strongly increased DOM concentrations and changed its characteristics. The magnitude of the observed changes was related to the upwelling intensity: more intense treatments led to higher accumulation of dissolved organic carbon (>70 μM of excess DOC over ambient waters for the most intense) and to comparatively stronger changes in DOM characteristics (increased proportions of chromophoric DOM (CDOM) and humic-like fluorescent DOM), suggesting a transformation of the DOM pool at the molecular level. Moreover, the single upwelling pulse resulted in higher CDOM quantities with higher molecular weight than the recurring upwelling mode. Together, our results indicate that under artificial upwelling, large DOM pools may accumulate in the surface ocean without being remineralised in the short-term. Possible reasons for this persistence could be a combination of the molecular diversification of DOM due to microbial reworking, nutrient limitation and reduced metabolic capabilities of the prokaryotic communities inside the mesocosms. Our study demonstrates the importance of the DOC pool when assessing the carbon sequestration potential of artificial upwelling.

## 1 Introduction

Primary producers and bacterioplankton require the uptake of inorganic nutrients from their surrounding waters to grow and keep their metabolism functioning (Azam and Malfatti, 2007). Inorganic nutrients are found in high concentrations below the photic layer (Johnson et al., 1997; Levitus et al., 1993), due to the absence of nutrient-consuming, light-driven primary production and the predominance of nutrient-releasing remineralization of organic matter. These nutrient-rich waters can reach the surface by a range of physical processes, such as winter convection (Severin et al., 2017), wind-driven coastal upwelling (Jacox et al., 2018), mesoscale eddies (McGillicuddy et al., 2007) or diapycnal diffusion (Arcos-Pulido et al., 2014), and play a key role in driving primary production and, consequently, CO_2_ fixation in the surface ocean (Field et al., 1998). A fraction of this newly produced organic matter is in turn exported, via multiple pathways, out of the photic layer into the deep ocean, where it is ultimately remineralised, releasing inorganic nutrients and carbon back to seawater. This process is known as the biological carbon pump (Le Moigne, 2019).

The biological carbon pump plays a major role in the atmosphere-ocean CO_2_ dynamics and acts as a key mechanism in the sequestration of CO_2_ in the deep ocean (Le Moigne, 2019). Nonetheless, there are great extents of the global oceans –such as the subtropical gyres that make up ∼40% of Earth’s surface (Polovina et al., 2008)– where the vertical input of nutrients is limited and, thus, phytoplankton rely on nutrients recycled within the photic layer, resulting in low primary production and CO_2_ fixation (Field et al., 1998). In the context of climate change and increasing CO_2_ emissions, the possibility of fertilizing these nutrient-poor waters to fuel primary production has been put forward as a climate intervention approach to enhance CO_2_ sequestration in the ocean and help reduce its atmospheric concentration (Fawzy et al., 2020; Williamson et al., 2012). One of the proposed fertilization methods consists in bringing nutrient-rich deep waters into the surface, an approach known as artificial upwelling (Pan et al., 2016). Artificial upwelling is envisioned to enhance the biological carbon pump by increasing primary production in nutrient-limited oceanic regions to amplify carbon export, ideally yielding a net increase in carbon sequestration if the C:N and C:P ratios of the exported organic matter are higher than the Redfield ratio. However, the efficiency of artificial upwelling to sequester atmospheric CO_2_ has been questioned (Shepherd et al., 2007; Yool et al., 2009), evidencing the need of further research.

Dissolved organic carbon (DOC) represents the largest pool of reduced carbon in the ocean and its downward flux is an important contributor to the biological carbon pump (Hansell et al., 2009; Le Moigne, 2019). The efficiency of the DOC pathway of carbon export will depend on the extent of its remineralization by prokaryotes and the depth at which it occurs in the water column. Some DOM is readily consumed by prokaryotes, and this process influences the absorption spectrum and fluorescence characteristics of DOM (Catalá et al., 2015b, 2018). A fraction of it however seems to escape remineralization and is accumulated. Two main hypothesis seek to explain this persistence (Dittmar et al., 2021): on the one hand, the intrinsic inability (or very reduced ability) of prokaryotes to consume some classes of organic molecules would lead to their accumulation and the long-term persistence of a fraction of DOM. On the contrary, the accumulated DOM might not be inherently resistant to degradation, its persistence being instead a consequence of complex ecological interactions between the vast molecular diversity of DOM (Zark et al., 2017b) and the wide metabolic capabilities of prokaryotes (Acinas et al., 2021; Sunagawa et al., 2015). In this scenario, all compounds would be continuously produced and degraded, stabilization occurring as a consequence of parallel decreases in the concentration of specific compounds and the abundance of prokaryotes able to degrade them (Dittmar et al., 2021). In summary, the balance between the transport of DOM to the ocean’s interior and how rapidly remineralization occurs above the permanent pycnocline (which depends on both physical dynamics and the ability of prokaryotes to consume DOM) will determine the net contribution of DOC to carbon sequestration. This balance will consequently influence the efficiency of artificial upwelling, making DOM a major factor to take into account when addressing carbon sequestration by this approach.

In the present work we studied the effect of the intensity and mode of artificial upwelling on the DOM pool in an oligotrophic marine environment. Using mesocosms that simulate ecological systems in close-to-natural conditions, we introduced nutrient-rich deep water into nutrient-depleted surface waters, aiming to investigate how the quantity, stoichiometry and composition of DOM is affected by upwelling and the subsequent enhancement of primary production. Deep-water addition was done simulating two modes of artificial upwelling: a singular addition, representing a moored system that upwells water into passing water patches, fertilizing them once (e.g., Zhang et al., 2016), and recurring deep water additions, simulating a system that drifts within a water parcel, repeatedly upwelling water into it (e.g., Maruyama et al., 2011). We aimed to study DOM dynamics under different artificial upwelling scenarios to gain insights into the short-term fate of DOM and the potential long-term implications for the efficiency of carbon sequestration.

## 2 Materials and methods

### 2.1 Experimental setup and sampling

Our experiment was conducted in Gando Bay (27° 55′ 41′′ N, 15° 21′ 55′′ W, Gran Canaria, Canary Islands) during the autumn of 2018 as a part of the Ocean artUp project. Nine KOSMOS (Kiel Off-Shore Mesocosms for Ocean Simulations; Riebesell et al., 2013) were deployed (M1-M9) and filled with in situ oligotrophic water (mean volume 43.775 ± 1.352 m^3^). To simulate artificial upwelling, nutrient-rich deep water was collected off Gran Canaria from 330 m (on day -10) and 280 m depth (on day 23), and added to the mesocosms in two different treatment modes (Table 1): a *singular* mode (M3, M7, M9, M1), in which a single deep-water (DW) addition was performed (on day 4), and a *recurring* mode (M2, M4, M6, M8), in which consecutive deep-water additions were performed (on days 4, 8, 12, 16, 21, 24, 28 and 32). For each artificial upwelling mode, four levels of intensity were simulated: *low* (M2, M3), *medium* (M7, M4), *high* (M6, M9) and *extreme* (M8, M1). Mesocosms within each intensity level had ultimately similar quantities of nutrients added to them (Table 1). A characterization of the DOM found in the deep water can be found in Table S1. No deep-water was added to M5 (*Control*) and ambient waters outside of the mesocosms were monitored during regular sampling (*Atlantic*). A detailed description of the experimental set up and sampling procedures can be found in Baumann et al. (2021b).

**Table 1.** Information of the treatments applied to each mesocosm. Total additions of deep water (as absolute values and % relative to the volume of the mesocosms), nitrogen (N), phosphorus (P) and silica (Si). N, P and Si values include both inorganic and organic forms.

| Mesocosm | Upwelling mode | Upwelling intensity | Added deep water volume [m<sup>3</sup>] | Added deep water volume [%]* | Added N [µmol·L<sup>-1</sup>] | Added P [µmol·L<sup>-1</sup>] | Added Si [µmol·L<sup>-1</sup>] |
| --- | --- | --- | --- | --- | --- | --- | --- |
| M5 | Control |  | 0.0 | 0.0 | 0.00 | 0.000 | 0.00 |
| M2 | Recurring | Low | 2.8 | 6.3 | 1.61 | 0.094 | 0.69 |
| M4 |  | Medium | 5.6 | 12.7 | 3.15 | 0.187 | 1.35 |
| M6 |  | High | 11.2 | 25.6 | 6.16 | 0.363 | 2.64 |
| M8 |  | Extreme | 22.4 | 51.1 | 10.97 | 0.682 | 4.96 |
| M3 | Singular | Low | 2.8 | 6.4 | 1.62 | 0.094 | 0.74 |
| M7 |  | Medium | 5.3 | 12.0 | 3.07 | 0.177 | 1.41 |
| M9 |  | High | 9.8 | 22.4 | 5.56 | 0.325 | 2.63 |
| M1 |  | Extreme | 17.2 | 39.2 | 9.80 | 0.567 | 4.58 |

\* Considering average mesocosm volume of 43.775 m<sup>3</sup>

Integrated samples of the water column within the mesocosms were collected using depth-integrated water samplers (IWS, HYDRO-BIOS, Kiel) and stored in acid-cleaned topaz bottles. To minimize photobleaching and degradation, samples were kept in dark, cool conditions until freezing/analysis (see below) on the same day. Prior to analysis/freezing, samples for dissolved organic matter quantification and characterization were filtered through precombusted (450ºC, 6h) glass fiber filters (GF/F, 0.7 μm nominal pore size) using acid-cleaned syringes and filter holders.

### 2.2 Dissolved Organic Carbon (DOC), Nitrogen (DON) and Phosphorus (DOP)

For DOC measurements, 10 mL of filtered samples were stored in high density polyethylene bottles and frozen (-20°C) until analysis. Samples were analyzed with a Shimadzu TOC-5000 analyzer (Sharp et al., 1993). Prior to the analysis, samples were thawed, acidified with 50 μL of H_3_PO_4_ (50%) and sparged with CO_2_-free air for several minutes to remove inorganic carbon. DOC concentrations were estimated based on standard curves (30–200 μM) of potassium hydrogen phthalate produced every day (Thomas et al., 1995). Reference material of deep-sea water (DSW, 42–45 μM C) provided by D. A. Hansell laboratory (University of Miami) was analyzed daily.

DON and DOP samples were collected in acid-rinsed, high-density polyethylene (HDPE) bottles. Upon arrival in the laboratory, 40 mL of the samples were filtered (0.45 μm cellulose acetate filters, Whatman) under sterile conditions. Total dissolved nitrogen and phosphorus were decomposed to phosphate and nitrate by adding an oxidizing solution and cooking the solution for approximately one hour. The samples were left to cool overnight, and the next day total dissolved nitrogen and phosphorus concentrations were measured spectrophotometrically on a continuous flow analyzer (QuAAtro AutoAnalyzer, SEAL Analytical). Triplicates of artificial seawater were treated and measured similarly on each measurement day. They acted as blanks and were averaged and subtracted from the samples. DON and DOP concentrations were calculated from the total dissolved nitrogen and phosphorus by subtracting the dissolved inorganic nitrogen and phosphate concentrations.

### 2.3 Chromophoric Dissolved Organic Matter (CDOM) characterization

Absorbance spectra were determined using an Ocean Optics USB2000+UV-VIS-ES Spectrometer alongside a World Precision Instruments liquid waveguide capillary cell (LWCC) with a path length of 0.9982 m. For each sample, absorbance was measured across a wavelength spectrum between 178 nm and 878 nm, performing a blank measurement prior to each sample using ultrapure Milli-Q water. Data processing was done in R (v. 4.1.2; R Core Team, 2021): raw spectra (samples and blanks) were cropped between 250 and 700 nm, blank spectra were subtracted from sample spectra and a dispersion correction was performed by subtracting the average absorbance of each sample between 600 and 700 nm to the whole spectrum.

After processing, absorbance was transformed into absorption following the definition of the Napierian absorption coefficient:

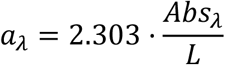

Where, for each wavelength λ, the absorption coefficient *a*_*λ*_ is given by the absorbance at wavelength λ (*Abs*_*λ*_), the path length of the cuvette (*L*, in meters; here 0.9982) and 2.303 (the factor that converts from decadic to natural logarithms).

From a_λ_ spectra values at 254 and 325 nm were considered. Both are proxies of CDOM concentration, although for different fractions of it: while a_254_ represents conjugated double bonds a_325_ is related to the aromatic fraction (Catalá et al., 2016, 2018; Lønborg and Álvarez-Salgado, 2014). Furthermore, spectral slopes between 275-295 nm and 350-400 nm were estimated from the natural log transformed absorption spectra following Helms et al. (2008). These regions of the spectra (as well as their ratio, S_R_) have been shown to be especially sensitive to changes in the molecular weight of CDOM, with higher slopes denoting lower average molecular weight (Helms et al., 2008, 2013). Moreover, these spectral slope parameters have been related to the microbial reworking of organic matter in the ocean, decreasing values being associated to increased transformation of DOM by prokaryotes (Catalá et al., 2015a, 2018).

### 2.4 Fluorescent Dissolved Organic Matter (FDOM) characterization

Fluorescence measurements were performed with a Jobin Yvon Horiba Fluoromax-4 spectrofluorometer, exciting the water samples in a wavelength range of 240–450 nm (10 nm increments), and measuring the fluorescence emission in a range of 300–560 nm (2 nm increments), with excitation and emission slit widths of 5 nm, and an integration time of 0.25 s. Fluorescence measurements were collected into excitation-emission matrices. To correct for lamp spectral properties and be able to compare results with other studies, excitation-emission matrices were measured in signal-to-reference mode with instrument-specific excitation and emission corrections applied during collection (Sc:Rc).

Excitation-emission matrices were processed using the DOMFluor toolbox (v. 1.7; Stedmon and Bro, 2008) for Matlab (R2017a). Alongside seawater samples, each sampling day three blank samples were measured using ultrapure Milli-Q water (at the beginning, middle and end of the measurement process). A weighted mean of the blanks was subtracted from each sample. Furthermore, excitation-emission matrices were normalized to the Raman area using the emission scan at 350 nm of ultrapure water blanks, calculating the area following the trapezoidal integration method (Lawaetz and Stedmon, 2009). Inner-filter correction was not performed as the average absorption coefficient of CDOM at 250 nm in all samples was 2.120 ± 0.544 m^−1^ (mean ± sd, n = 208; max. = 3.4 m^−1^), which was lower than the threshold of 10 m^−1^ above which this correction is considered to be necessary (Stedmon and Bro, 2008). Rayleigh scatter bands of 1st (*Em = Ex ± bandwidth*) and 2nd (*Em = 2·Ex ± 2·bandwith*) orders were cut at each wavelength pair.

### 2.5 Parallel Factor analysis (PARAFAC) of fluorescence data

The processed excitation-emission matrices (n = 175, samples with measurement errors were removed) were analyzed applying a PARAFAC analysis (Stedmon et al., 2003; Stedmon and Bro, 2008) using the DOMFluor toolbox. A model consisting of five components (named according to their emission maxima; Table S2, Figure S1) was validated by split-half validation and random initialization. For each sample, the fluorescence maximum (F_max_) of the components was recorded.

The optical characteristics of the five components are summarized in Table S2, along with similar fluorophores found in the literature. The identification of previously described fluorophores was performed using the OpenFluor database (openfluor.lablicate.com, Murphy et al., 2014), based on the combined Tucker Congruence Coefficient of the excitation and emission spectra (TCC_ex·em_). C1, C2, C4 and C5 had 17, 28, 11 and 6 matches with high congruence (TCC_ex·em_ >0.95), respectively. C1 presented characteristics similar to fluorophores identified as amino acid-/tryptophan-like, with primary and secondary excitation maxima at 300 and 240 nm, respectively, and an emission maximum at 354 nm (Table S2). Such amino acid-like compounds have been previously described as partially bioavailable for prokaryotic consumption (Lønborg et al., 2010). C2 (excitation maxima at 250 and 330 nm, emission maximum at 410 nm) displayed high similarities with humic-like fluorophores (peak M, Table S2) that have been observed to be positively correlated to apparent oxygen utilization in the ocean (Catalá et al., 2015b). C4 was also similar to previously identified humic-like components but, unlike C2, its signal presented peaks at higher wavelengths (excitation maxima at 260 and 370 nm, emission maximum at 466 nm). It resembled a mixture of peaks A and C, formed by compounds with high aromaticity (Table S2). Similarly to C1, C5 also presented excitation and emission spectra highly congruent with amino acid-/tryptophan-like fluorophores, but had lower maxima than C1 (excitation and emission maxima at 270 and 342 nm, respectively; Table S2). Notably, the spectrum of this component was highly similar to that of indole (Wünsch et al., 2015). As opposed to the other components, C3 presented matches with congruence lower than 0.95. With excitation maximum below 240 nm and a broad emission spectrum (maximum at 330 – 472 nm), it was identified as similar to fluorophores potentially related to fluorometer artifacts (Table S2). Thus, the C3 component was not further considered in the analyses.

### 2.6 Prokaryotic heterotrophic production and cell abundance

Prokaryotic heterotrophic production (PHP) was estimated via the incorporation of ^3^H-leucine using the centrifugation method (Smith and Azam, 1992). ^3^H-leucine (Perkin-Elmer, specific activity 160 Ci mmoL^-1^) was added at final concentration (20 nmol L^-1^) to quadruplicate 1 mL subsamples. Blanks were established by adding 100 μL of 50% trichloroacetic acid (TCA) to duplicate blanks screw-cap microcentrifuge tubes 15 min prior to radioisotope addition. The microcentrifuge tubes were incubated at *in situ* temperature (± 1ºC) in the dark for 2 h. Incorporation of leucine in the quadruplicate tubes was stopped by adding 100 μL ice-cold 50% TCA and tubes were kept together with the blanks at 4ºC until further processing as in Smith and Azam (1992). The mean disintegrations per minute (DPM) of the TCA-killed blanks were removed from the mean DPM of the respective samples and succeeding DPM value converted into leucine incorporation rates. PHP was calculated using a conservative theoretical conversion factor of 1.55 kg C moL^-1^ Leu assuming no internal isotope dilution (Kirchman, 1993). The PHP data is available at the PANGAEA repository (Baumann et al., 2021a).

Samples for prokaryotic cell abundance were collected into 2 mL cryovials, fixed with a 2% final concentration of paraformaldehyde and stored at -80°C. After thawing, 400 μL subsamples were stained with 4 μL of the fluorochrome SYBR Green I (Molecular Probes) diluted in dimethyl sulfoxide (1:10) and analyzed in a FACSCalibur (Becton-Dickinson) flow cytometer. Fluorescent beads (1 μm, Polysciences) were added for internal calibration (10^5^ mL^-1^). Prokaryotic cells were identified in green fluorescence vs side scatter cytograms.

### 2.7 Inorganic nutrients, Chlorophyll-a and particulate organic carbon

Nitrate (NO_3_^−^), nitrite (NO_2_^−^), ammonium (NH_4_ ^+^), phosphate (PO_4_^3−^) and silicic acid (Si(OH)_4_) were quantified spectrophotometrically on a five channel continuous flow analyzer (QuAAtro AutoAnalyzer, SEAL Analytical Inc., Mequon, United States). Chlorophyll-a (Chl-a) was measured with an HPLC Ultimate 3,000 (Thermo Scientific GmbH, Schwerte, Germany). Particulate organic carbon (POC) in the water column was measured using a CN analyzer (Euro EA-CN, HEKAtech). See Baumann et al. (2021b) for details. Chl-a and POC data are available at the PANGAEA repository (Baumann et al., 2021a).

### 2.8 Statistical analyses

All statistical analyses and data representations were performed in R (v. 4.1.2; R Core Team, 2021). Linear regressions were performed to assess the relationships between upwelling intensity (as N addition, in μM) and the dissolved organic matter variables. Normality and homoscedasticity of residuals were tested with Shapiro-Wilk (*stats* package, v. 4.1.2) and Breusch-Pagan (*lmtest* package, v. 0.9.39; Zeileis and Hothorn, 2002) tests, respectively. Data representations were done with *ggplot2* (v. 3.3.5; Wickham, 2016).

## 3 Results

### 3.1 Response of primary producers

Artificial upwelling led to large phytoplankton blooms after the first deep water addition, increasing primary production and shifting the community composition towards a diatom dominated assemblage (Ortiz et al., 2022). Changes were in accordance with the intensity level of upwelling: the largest Chl-a build-ups were registered in the singular extreme treatment, were Chl-a reached 11.2 μg·L^-1^ on day 9 (Fig. 1A). Upwelling modes however differed in their outcomes after the first deep water addition. In the singular treatments the blooms rapidly collapsed and Chl-a remained low until the end of the experiment, while in the recurring treatments subsequent deep water additions allowed to maintain the blooms (with oscillating Chl-a concentrations, Fig. 1A) throughout the experiment. Particulate organic carbon (POC) concentrations in the water column (Fig. 1B) showed accumulations following the phytoplankton bloom dynamics. Singular treatments accumulated the highest POC values after the initial bloom, reaching 66 μM in the extreme intensity level, followed by a steady decrease. The response in the recurring treatments was slower but POC accumulated until day ∼20 (63 μM in the extreme treatment). While the least intense levels ended with similar concentrations to those of the singular mode, in the high and extreme recurring treatments POC values markedly a after day 27 (maximum of 102 μM) linked to coccolithophorid blooms at the end of the experiment (Ortiz et al., 2022).

**Figure 1.**
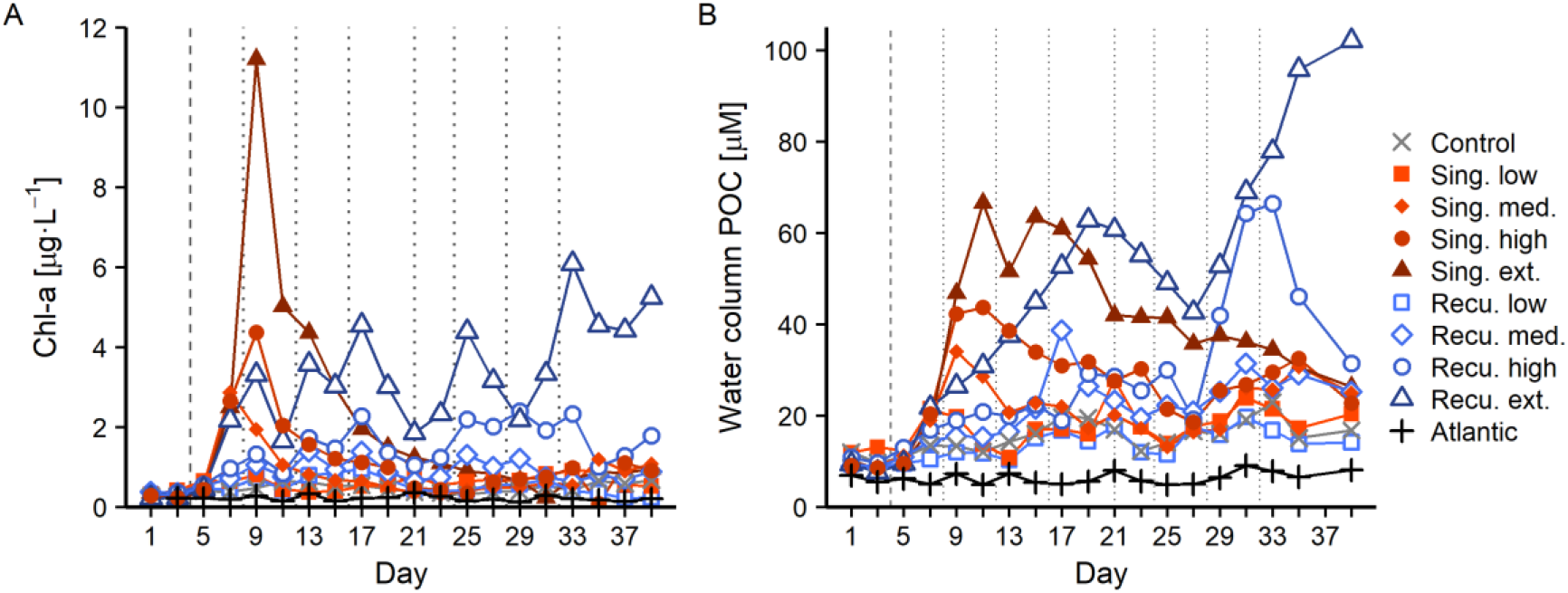
Chl-a and particulate organic carbon (POC) concentrations during the experiment. Vertical lines indicate deep water additions of singular (dashed) and recurring (dashed and dotted) treatments.

### 3.2 Dissolved organic matter concentration and elemental composition

Artificial upwelling yielded large increases in the DOM pool, in direct positive relationship with upwelling intensity (Figure 2 and 3). Extreme treatments presented DOC concentrations that were +70 μM compared to starting conditions (Figure 2A). Average DOC concentrations prior to the first deep water addition at day 4 ranged between 70.6 – 78.1 μM. After the deep water addition DOC remained relatively stable without exceeding initial values until day 9, when concentrations started to raise in all treatments coinciding with the peak of the diatom bloom (Figure 1). After day 9, while the bloom in the singular treatments started to collapse, DOC concentrations continued to increase, especially in the singular treatments: on day 13, the extreme, high and medium singular treatments showed the highest DOC values with 155, 109 and 106 μM respectively (Figure 2A). After day 13, DOC in these mesocosms did not continue to increase and tended to stabilize until the end of the experiment.

**Figure 2.**
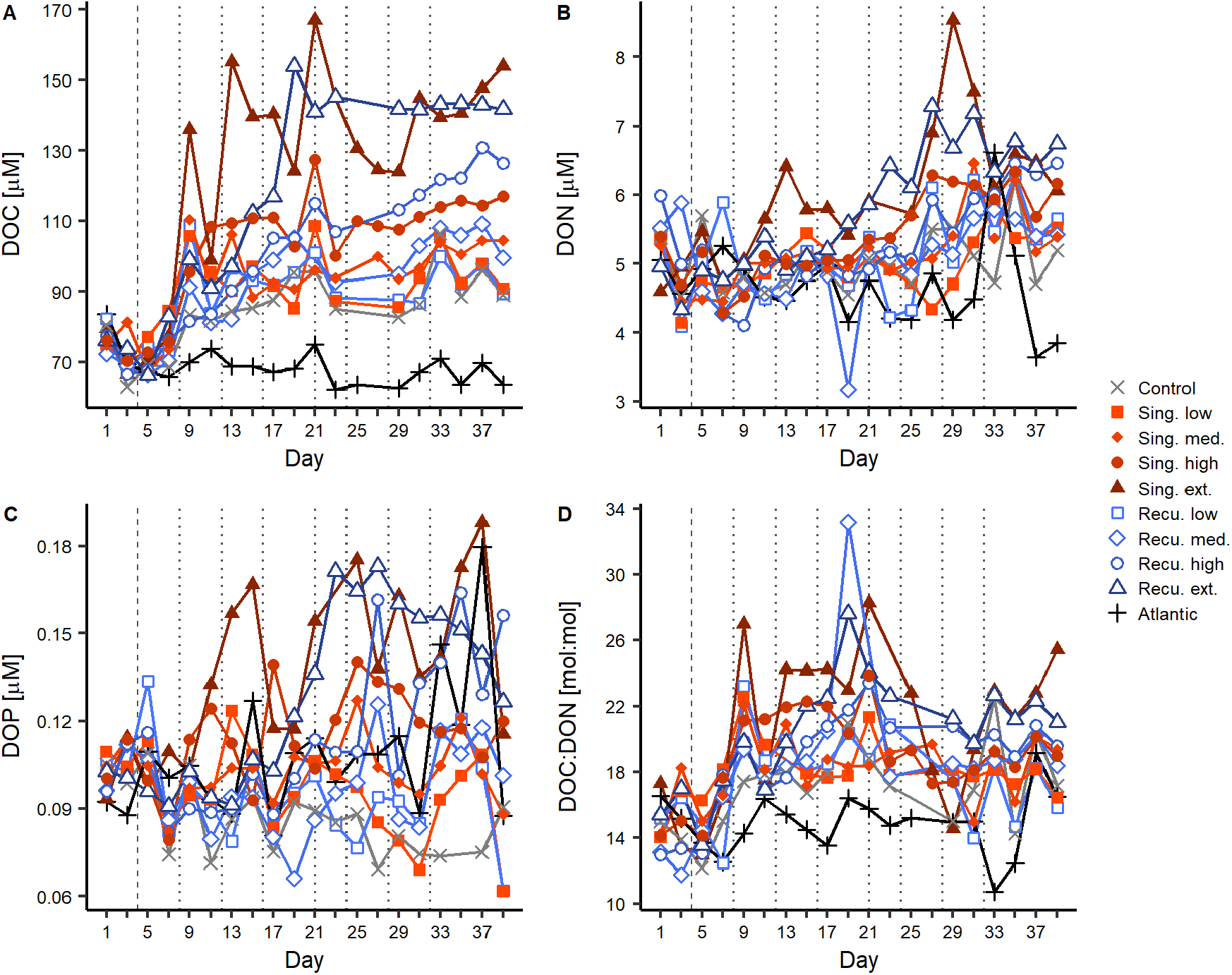
Changes in dissolved organic matter concentrations and ratios during the experiment. Temporal evolution of A) DOC, B) DON, C) DOP, D) DOC:DON. Vertical lines indicate deep water additions of singular (dashed, day 4) and recurring (dashed and dotted) treatments. Results for DOC:DOP and DON:DOP ratios can be found in Figures S1.

**Figure 3.**
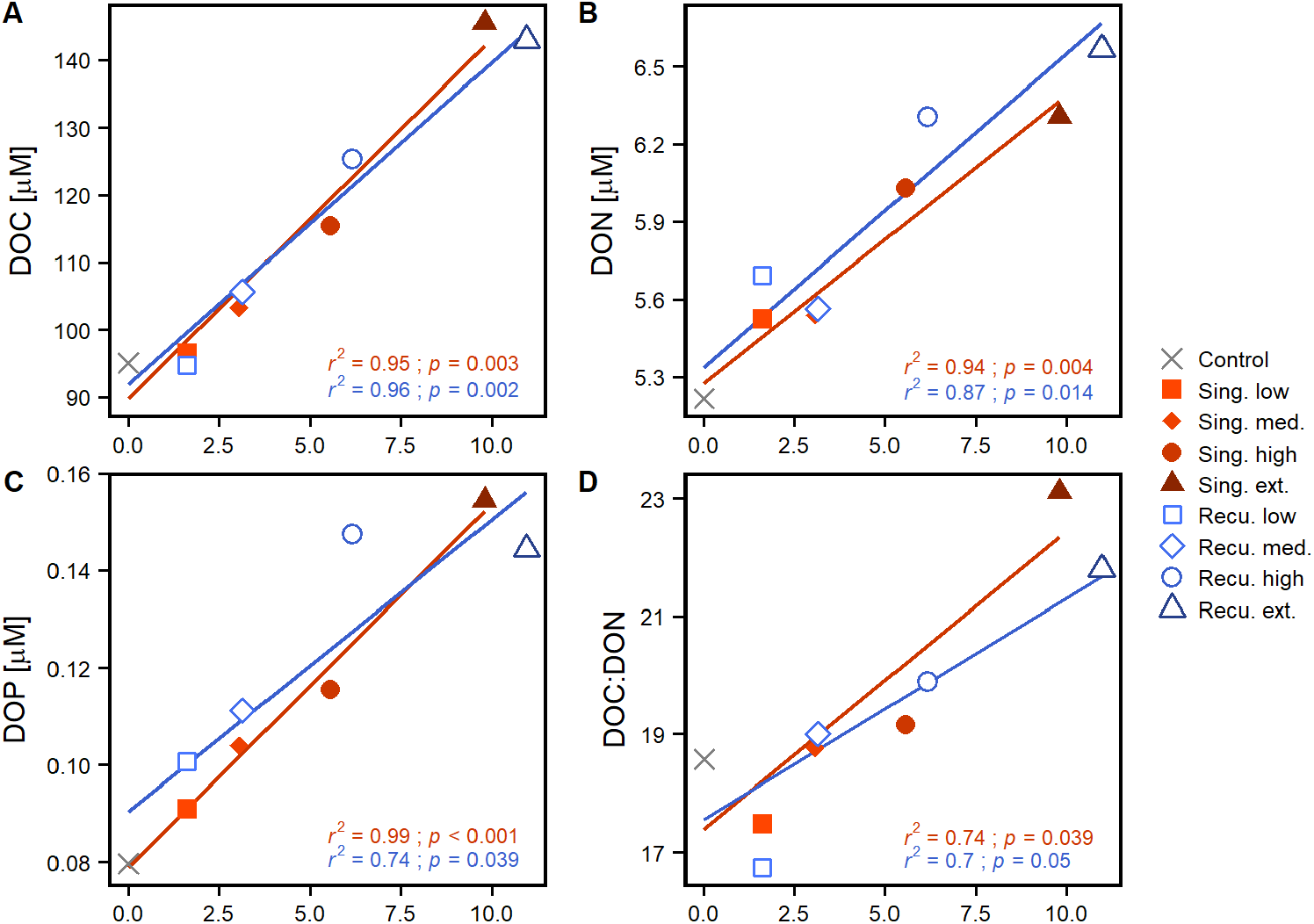
Effect of upwelling intensity on dissolved organic matter. Shown are linear regressions of A) DOC, B) DON, C) DOP and D) DOC:DON values against upwelling intensity (N addition) per upwelling mode. DOM values were averaged after the last deep water addition (≥ day 33). The coefficient of determination (*r*^*2*^) and p-value (*p*) of the regressions are included. Only the lines for significant regressions (*p* < 0.05) are displayed (see Table S3 for detailed test statistics). Results for DOC:DOP and DON:DOP ratios can be found in Figure S2.

In the recurring treatments, where phytoplankton abundances experienced smaller initial increases but did not collapse (Figure 1), DOC increases were not as abrupt. Only between days 16-20, when the bloom in the extreme recurring treatment presented a major decrease, DOC displayed pronounced increases, reaching its peak on day 19 (154 μM) and subsequently sustained similar values, while the high treatment experienced a steady increase throughout the experiment (Figure 2A). During days 33-39, average DOC values were of 143-145, 115-125, 103-106 and 95-97 μM for extreme, high, medium and low treatments, respectively. After the final deep water addition to the recurring treatments (days ≥33), average DOC concentrations showed a significant positive relationship with upwelling intensity (Figure 3A), with a similar effect for both upwelling modes (Table S3). During the entire experiment, DOC concentrations in the control mesocosm trailed those in the low treatments, while the Atlantic remained relatively stable with far smaller values (mean ± sd = 68.5 ± 5.1 μM).

Increases in DON and DOP concentrations were more subtle than in DOC (Figure 2B and 2C). Initial values ranged between 4.65-5.70 μM and 0.102-0.109 μM, respectively. At day 11-13 DON values started to raise in the extreme singular treatment, followed by the extreme recurring treatment at day 19. These mesocosms reached maximum values of 7-8 μM during days 27-31. In contrast to DOC concentrations, DON values departed less from initial conditions and presented higher variability but increasing DON concentrations were associated with increasing upwelling intensity, with similar effects for both modes (Figure 2B and 3B, Table S3). Similarly, differences in DOP between treatments were minor until day 11, when the extreme singular treatment started to increase, reaching values of >0.15 μM and fluctuating until the end of the experiment. The extreme recurring treatment began to separate from less intense treatments in day 19, peaking during days 23-27 at ∼0.17 μM and slowly decreasing towards the end (Figure 2C). Despite variability, overall treatments subject to more intense upwelling presented higher DOP concentrations: average DOP values in days ≥33 showed significant positive relationships with upwelling intensity both for singular and recurring upwelling modes (Figure 3C, Table S3).

The differential changes in DOC, DON and DOP were reflected in the elemental ratios (Figure 2D and S2), with notable increases in the values of the DOC:DON ratio, leading to C-enriched DOM. The average DOC:DON:DOP ratio across mesocosms prior to the first deep water addition was 704:48:1 (DOC:DON ∼15:1). After the addition and following the DOC build up, carbon ratios started to increase, although consistent differences between treatments were only observed for DOC:DON (Figure 2D), but not DOC:DOP (Figure S2). During days 13-21, DOC:DON ratios were highest, particularly in mesocosms with more intense upwelling: extreme singular and recurring treatments showed average values of 25:1 and 23:1, respectively, while low treatments only reached 19:1. Average DOC:DOP ratios ranged between 927:1 – 1178:1 for this same period. During days 33-39, DOC:DON values experienced a slight decrease, but still were higher than initial ratios and showed consistent differences between treatments (17:1 – 23:1 across intensity levels). In fact, DOC:DON showed a positive relationship with upwelling intensity (Figure 3D).. While DON:DOP values showed considerable variability and no consistent temporal trend throughout the experiment (Figure S2), the more intense treatments tended to show lower values. After the last deep water addition, DON:DOP ratios showed negative relationships with upwelling intensity, although only significant for the singular treatments (Figure S3).

### 3.3 Optical characteristics of dissolved organic matter

The chromophoric and fluorescent fractions of the dissolved organic matter presented pronounced increases in response to artificial upwelling. CDOM concentrations, as depicted by a_254_ (Figure 4A), increased in all mesocosms once the first water addition was done (including the control despite no nutrients were added). Initial values ranged between 1.22-1.39 m^-1^ and saw a steady increase for most of the experiment, reaching 2.90 m^-1^ (extreme recurring) and 3.17 m^-1^ (extreme singular). The extreme and high singular treatments displayed a pronounced increase on days 9-17 (during and post phytoplankton bloom, Figure 1), and the extreme recurring treatment on days 13-17, but subsequently continued to raise at a gentler rate until the end of experiment. Resulting CDOM quantities were higher in mesocosms with more intense simulated upwelling and were overall greater in the singular treatments: recurring treatments displayed values that were comparable to the previous intensity level in singular treatments (Figure 4A). While both upwelling modes showed significant positive relationships between a_254_ and upwelling intensity (Figure 5A), the slope of the singular mode was steeper than the recurring one even when considering the 95% confidence interval (Table S3). The aromatic fraction of CDOM, represented by a_325_ (Figure S4), followed patterns very similar to a_254_, starting at 0.17-0.19 m^-1^ and ending at 0.53-0.99 m^-1^, and also showed a positive relationship with upwelling intensity (Figure S5, Table S3).

**Figure 4.**
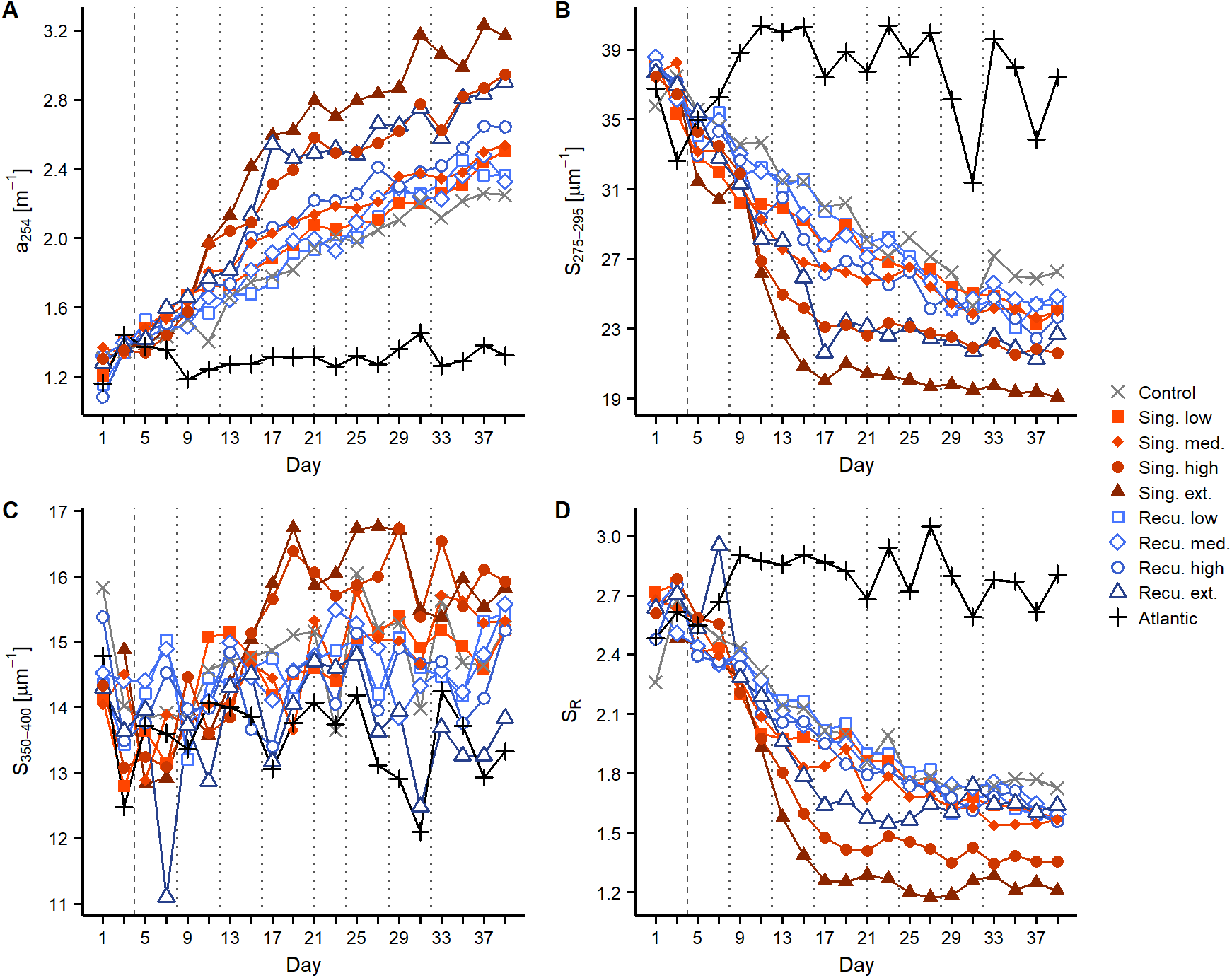
Changes in chromophoric dissolved organic matter during the experiment. Temporal development of a) CDOM quantity as a_254_, b) S_275-295_, c) S_350-400_, and d) SR. Vertical lines indicate deep water additions of singular (dashed) and recurring (dashed and dotted) treatments. See explanation of parameters in the Methods section.

**Figure 5.**
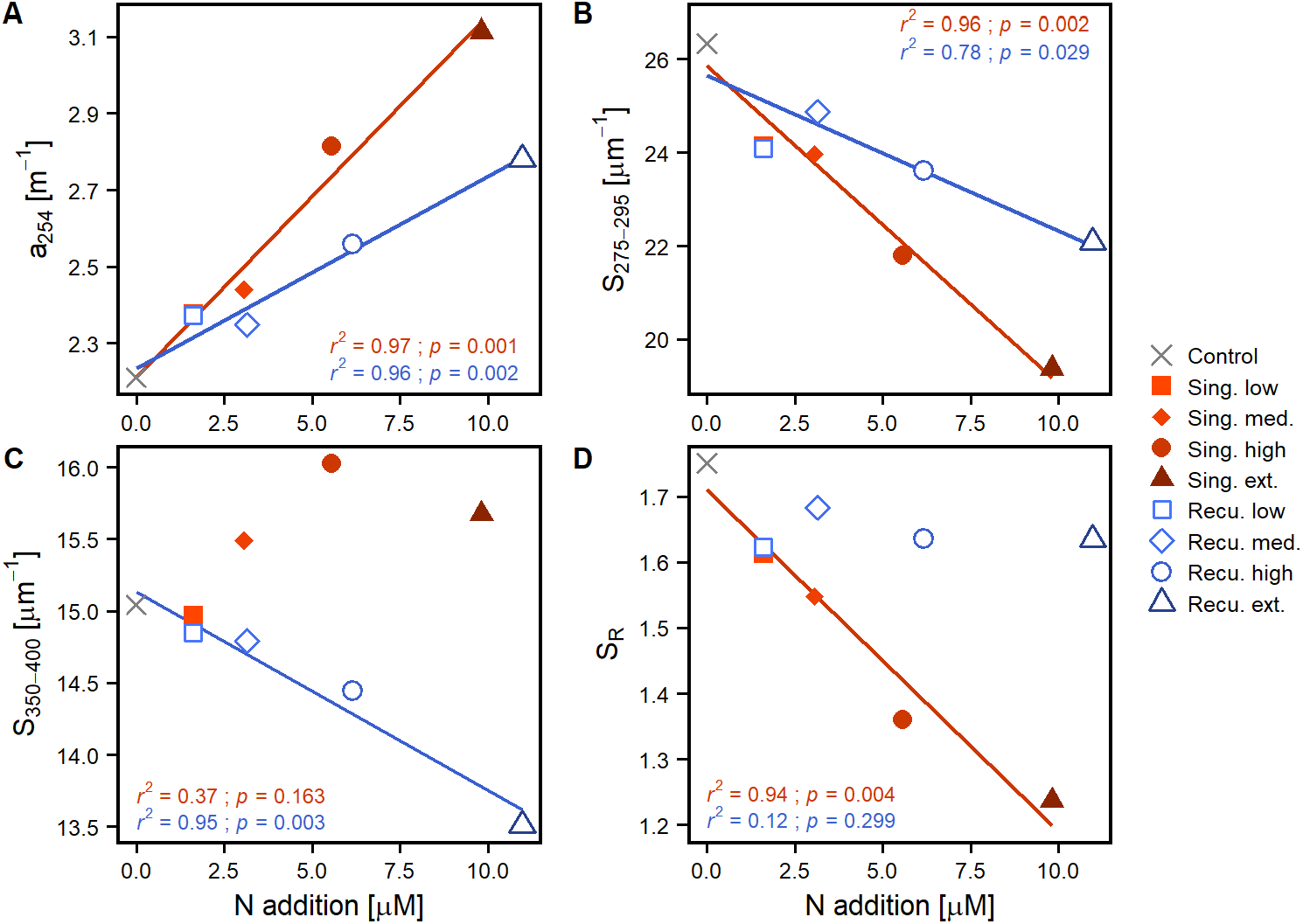
Linear regressions of average values after the last deep water addition to recurring treatments (≥ day 33) of A) CDOM quantity as a_254_, B) S_275-295_, C) S_350-400_ and D) S_R_ against upwelling intensity (as N addition), per upwelling mode. The coefficient of determination (*r*^*2*^) and p-value (*p*) of the regressions are included. Only lines for significant regressions (*p* < 0.05) are displayed. Regression parameters are detailed in Table S3.

Spectral slopes (S_275-295_ and S_350-400_), which provide insights into the average molecular weight of the CDOM pool, also experienced marked changes. S_275-295_ (Figure 4B) began at 36.0-37.9 μm^-1^ and decreased throughout the experiment, signaling an increase in average molecular weight, as compounds of higher molecular weight tend to absorb light at higher wavelengths, thus decreasing the slopes. Coupled with changes in a_254_, S_275-295_ in the high and extreme singular treatments showed a very pronounced decline between days 9-13, while a sharp decrease was registered on days 13-17 in the extreme recurring treatment. After day 17 values tended to stabilize in the more intense simulated upwelling treatments while less intense treatments continued to steadily decrease (Figure 4B). The extreme singular treatment displayed the lowest S_275-295_, reaching average values of 19.4 ± 0.3 μm^-1^ during days ≥33. Average S_275-295_ values for this period showed significant negative relationship with upwelling intensity for both modes (Figure 5B) but, as for a_254_, S_275-295_ values in singular treatments were comparable to the previous intensity level in recurring treatments. S_350-400_ (initial values of 13.5-14.9 μm^-1^, Figure 4C) on the other hand presented opposite trends for each upwelling mode: while singular treatments presented increases (the extreme reaching 16-17 μm^-1^ on days 17-29), the recurring treatments tended to decrease. This resulted in different outcomes after day 33: despite the increase, singular treatments did not show a significant relationship with upwelling intensity, while recurring treatments showed a significant negative one (Figure 5C). Resulting S_R_ values overall were dominated by the marked changes in S_275-295_ and followed its patterns (Figure 4D): initially at 2.48-2.74, decreases were observed specially in high and extreme singular patterns (the latter reaching minimum values close to 1.25) until day 17 and subsequently tended to stabilize. However, the different fates of S_350-400_ for the two upwelling modes were reflected in S_R_: at the end of the experiment, singular treatments displayed a significant relationship with upwelling intensity (Figure 5D), but recurring ones did not and tended to converge around S_R_ values of 1.65 (Figure 4D).

Fluorescence measurements provided further details into the composition of the DOM pool. Components C1 and C5 of the PARAFAC model were similar to fluorophores described as amino acid-like/tryptophan-like compounds that are at least partially bioavailable to prokaryotic consumption (Table S2). Both C1 and C5 (Figures 6A and 6D) presented increases in the intensity of their signal across mesocosms after the first deep water addition. For C1 the extreme singular treatment experienced a fluorescence intensification that was clearly superior to any other treatment, increasing from 0.009 RU to 0.070 RU in day 15, and continued to increase until reaching 0.087 RU at the end of the experiment. Other treatments also displayed increases until day 15 (although smaller) and tended to stabilize afterwards, despite variability, ending within a range of 0.032 (high singular) and 0.044 RU (low recurring). Despite the extreme singular treatment showed markedly high values, no consistent relationship was found between C1 and upwelling intensity at the end of the experiment (Figure 7A). For C5 (Figure 6D) all treatments exhibited relatively similar patterns. Initial values ranged between 0.001-0.007 RU, fluorescence signals increasing until day 21 and subsequently maintaining relatively similar values, ending at 0.011-0.021 RU. C5 did display significant positive relationships with upwelling intensity for both upwelling modes (Figure 7D).

**Figure 6.**
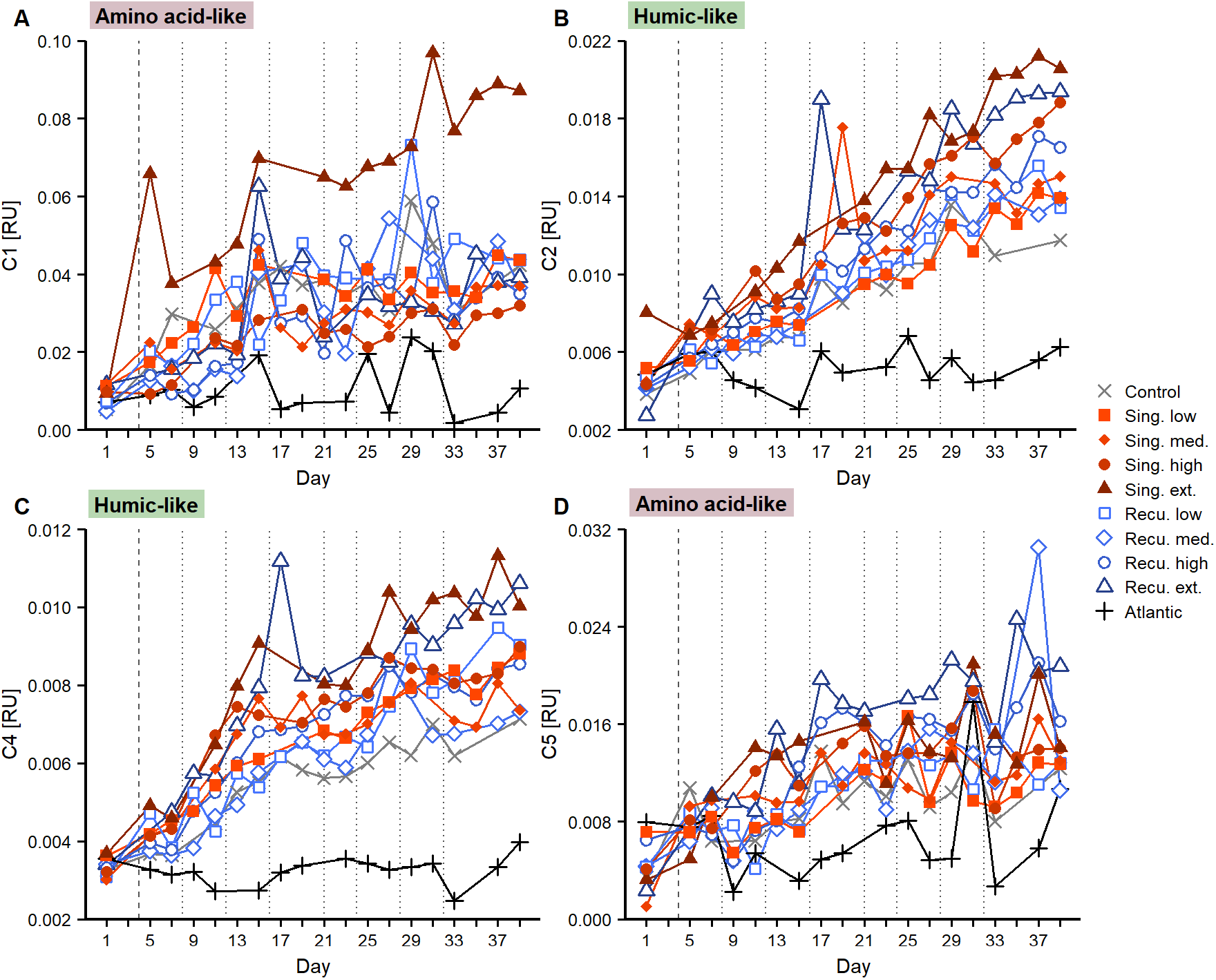
Changes in fluorescent dissolved organic matter during the experiment. Temporal evolution of PARAFAC components A) C1, B) C2, C) C4 and D) C5. Vertical lines indicate deep water additions of singular (dashed) and recurring (dashed and dotted) treatments.

**Figure 7.**
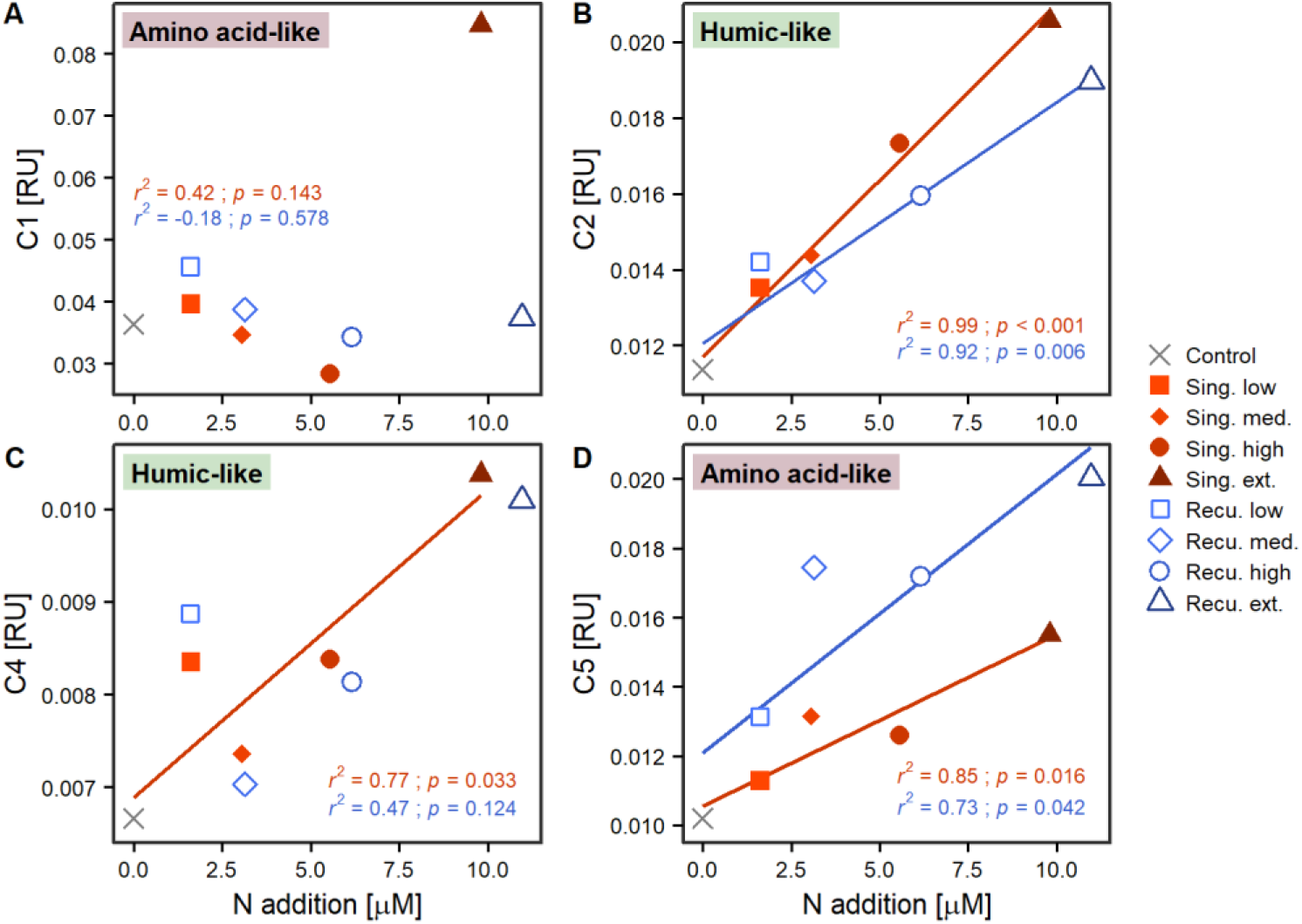
Linear regressions of average values after the last deep water addition to recurring treatments (≥ day 33) of FDOM components A) C1, B) C2, C) C4 and D) C5 against upwelling intensity (as N addition), per upwelling mode. The coefficient of determination (*r*^*2*^) and p-value (*p*) of the regressions are included. Only lines for significant regressions (*p* < 0.05) are displayed. Regression parameters are detailed in Table S3.

Components representing humic-like compounds (C2 and C4), which have been associated to the microbial transformation of DOM, displayed nearly continuous increases in fluorescence signals (Figure 6B and C). C2 started at values of 0.003-0.008 RU and increased throughout the experiment in all mesocosms, treatments with greater upwelling intensity exhibiting more intense fluorescence signals. Positive significant relationships were found with upwelling intensity after the final deep water addition (Figure 7B). Singular treatments presented slightly higher values than recurring ones: e.g., during days ≥33, average C2 values for the extreme singular and recurring treatments were 0.0206 ± 0.0005 and 0.0190 ± 0.0005 RU, respectively. C4 also displayed fluorescence signals that consistently increased throughout the experiment (Figure 6C) and, as C2, presented significant positive relationships with upwelling intensity for both modes at the end of the experiment (Figure 7C).

## 4 Discussion

### 4.1 DOC accumulation in the water column

Initial concentrations of DOC were similar to those typically found in surface waters of the Canary Islands oceanic region (Arístegui et al., 2003, 2004, 2020; Burgoa et al., 2020) and other locations in the North Atlantic subtropical gyre (Goldberg et al., 2010; Lomas et al., 2013). Concentrations quickly increased following nutrient depletion and the collapse of the diatom-dominated phytoplankton bloom (Ortiz et al., 2022). In the case of extreme treatments (∼140 μM), values reached levels far exceeding DOC concentrations observed in the Canary Current upwelling system (100-110 uM, Arístegui et al., 2003; Burgoa et al., 2020). Similar DOC levels have previously been reported in deep water addition mesocosm experiments in oligotrophic waters (Zark et al., 2017a).

Diatoms are known to release DOC upon nutrient limitation (Norrman et al., 1995; Zark et al., 2017a), but the source of the observed DOC (∼70 μM increase in extreme treatments) was not limited to dissolved primary production. Accumulated dissolved primary production by day 21 was 20.53 μmol C·L^-1^ (11.91% of total primary production) and 6.58 μmol C·L^-1^ (5.07% of total primary production) for the extreme singular and recurring treatments, respectively (Figure S6; Ortiz et al., 2022). Given these values, most of the DOC increase must have originated from a source other than dissolved primary production. Considering the high accumulated particulate primary production, values (Figure S6) and that more than a half of it was retained in the water column during the experiment (Baumann et al., 2021b), the channeling of carbon from the particulate to the dissolved fraction likely accounted for the rest of the DOC increase.

The arising question is how such vast amounts of organic carbon were channeled into the dissolved matter pool in such a short period of time (∼10 days). Processes such as extracellular enzymatic activity on particles and gel structures (Arnosti, 2011), grazing and sloppy feeding (Steinberg and Landry, 2017) or viral lysis (Breitbart et al., 2018) result in release of DOM. The large amounts of particles (including transparent exopolymer particles) that were formed following the diatom bloom (Baumann et al., 2021b) provided the substrate for prokaryotes to consume POC, and this could have contributed to the production of DOC. Moreover, viral lysis can be key to phytoplankton bloom termination, resulting in major releases of DOC (Breitbart et al., 2018; Highfield et al., 2014). The balance of DOC after day 21 despite continued primary production, in conjunction with sustained community respiration (Baños et al., in prep.) and prokaryotic heterotrophic production (PHP) rates (Figure S7), indicates that prokaryotes were consuming at least a fraction of the DOM pool and, consequently, at least part of the DOC accumulated in the water column could eventually be remineralised.

### 4.2 Carbon enrichment in DOM

Artificial upwelling enhanced the C:N ratio in the DOM pool from ∼15 to 19-27. Initial DOC:DON values (Figure 2D) were within the range of values reported for the subtropical North Atlantic (Hansell and Carlson, 2001; Valiente et al., 2022). Subsequent increases in DOC:DON ratios as a result of artificial upwelling were similar to those observed in other experimentally-induced diatom blooms (Norrman et al., 1995). This increase was probably a combination of the release of C-rich polysaccharides by diatoms (Engel, 2001; Mühlenbruch et al., 2018) and the preferential degradation of DON by prokaryotes (Hach et al., 2020). On the contrary, no clear relationship with upwelling intensity was observed for DOC:DOP (Figure S2 and S3), although overall values slightly increased from initial conditions and, thus, preferential DOP consumption might have existed to a certain degree (Hach et al., 2020). While DOP concentrations fluctuated considerably (Figure 2C) overall DON:DOP values decreased with increasing upwelling intensity at the end of the experiment (Figure S2). Considerable variability exists in C:N:P ratios of heterotrophic prokaryotes, with taxa presenting N:P ratios both below (White et al., 2019) and similar to (Zimmerman et al., 2014) Redfield values. However, given that nutrients were added close to Redfield proportions (Table 1), but NO_3_^−^:PO_4_^−^ ratios decreased after deep water additions (Figure S8), DON might have been consumed over DOP to compensate for the lower NO_3_^−^ availability.

### 4.3 Shift towards high molecular weight, humic-like DOM

CDOM increases have been previously observed associated with phytoplankton production (Romera-Castillo et al., 2010), including communities in nutrient-depleted conditions after a blooming phase (Loginova et al., 2015). Hence, as with DOC, the initial post-bloom increase in a_254_ (Figure 4A) and a_325_ (Figure S4) was probably partly associated with DOM released by diatoms. While DOC tended to stabilize, CDOM continued to accumulate, including an intensification of the humic-like fluorescence signal (Figure 6). The sustained accumulation throughout the experiment seems to be the result of the generation of CDOM and humic-like FDOM as by-products of the prokaryotic reworking of organic matter (Catalá et al., 2015b; Nelson and Siegel, 2013). The fact that cumulative PHP was strongly correlated with CDOM absorption, spectral slopes and humic-like FDOM intensity (Figure 8) would support that interpretation, as cumulative PHP has been previously observed to be correlated to DOM transformation (Zark et al., 2017a).

**Figure 8.**
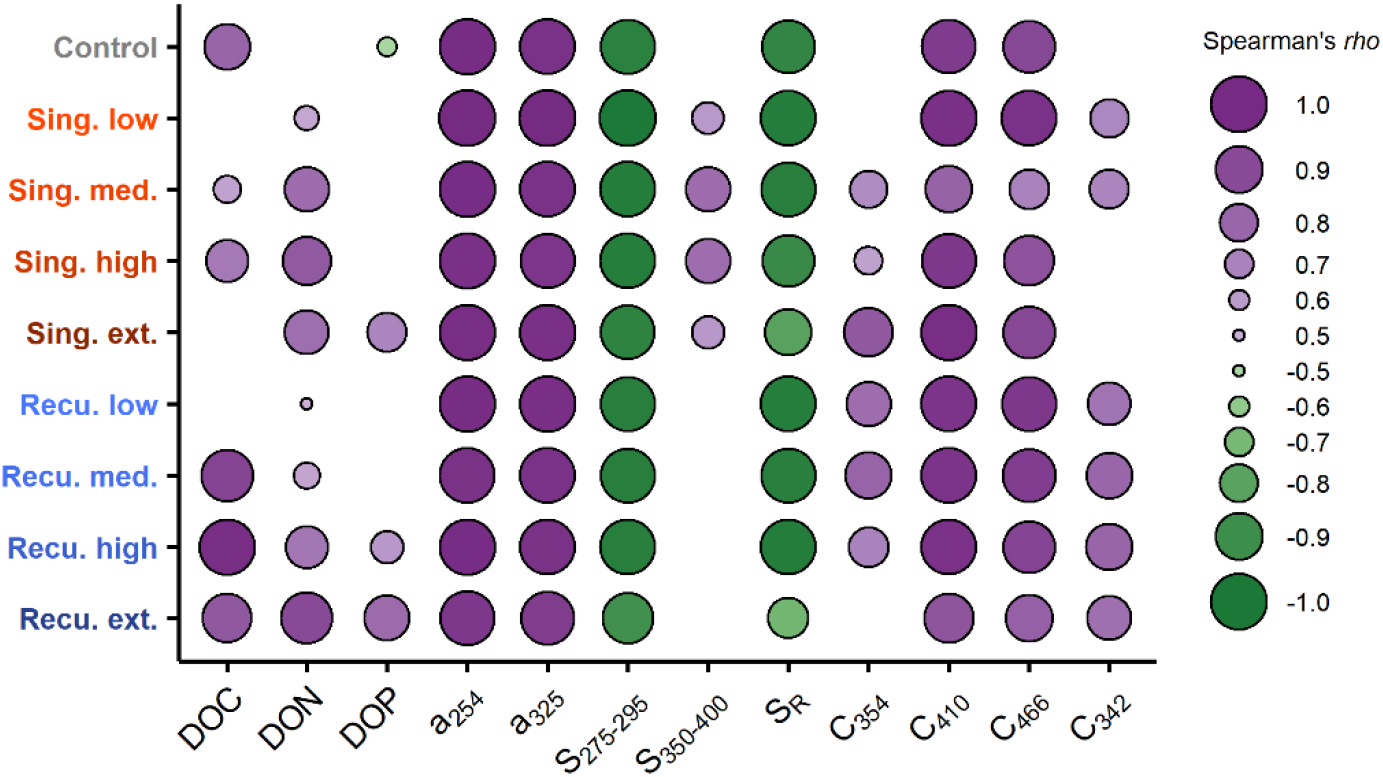
Relationship between cumulative prokaryotic heterotrophic production (PHP) and dissolved organic matter (DOM) parameters. Shown are Spearman’s *rho* (*ρ*) correlation coefficients per mesocosm. Only significant correlations (p < 0.05) are displayed.

Diatoms are also known to release amino acids (Granum et al., 2002) and the initial increase in amino acid-like fluorescence (Figure 6) was probably an indication of the release of molecules containing amino acids (e.g. free amino acids, proteins and aminosugars) by the blooming community. This group of molecules represents one of the most labile fractions of DOM and are known to undergo rapid biodegradation (Benner and Amon, 2015). Over the course of the experiment the amino acid-like fluorescence was not clearly related to upwelling intensity, which may indicate ongoing consumption by prokaryotes. However, values did not return to initial levels, meaning that a fraction of this signal represented molecules that were not readily consumed by prokaryotes. In DOM degradation experiments carried out in a coastal upwelling system, Lønborg et al. (2010) reported a similar amino acid-like fluorescence remnant that they attributed to a fraction of it being biologically unavailable. Amino acid release through viral lysis (Middelboe and Jørgensen, 2006) and, for recurring treatments, more constant releases of amino acid-containing molecules by the prolonged diatom blooms throughout the experiment (Ortiz et al., 2022) could have also contributed to the sustained amino acid-like fluorescence signal.

CDOM average molecular weight also changed markedly: initial values of spectral slopes were similar to those found in other oligotrophic regions (e.g. Catalá et al., 2018) but markedly decreased after the initial bloom (Figure 4B-C), suggesting an increase in average molecular weight (Helms et al., 2008). Artificial upwelling modes, however, had different outcomes, with higher CDOM molecular weight in the singular treatments and differences in composition (Figs. 4B-D). According to the size-reactivity continuum (SRC) theory (Benner and Amon, 2015), larger size classes of DOM (e.g., macromolecules of combined forms of carbohydrates and amino acids) tend to be preferentially degraded by prokaryotes. The degradation of this high molecular weight DOM, which here probably included polysaccharides and molecules containing amino acids, would result in the generation of CDOM (and FDOM) of higher molecular weight than what was initially present in the mesocosms, hence yielding the observed changes in spectral slopes. Similar results have been reported for the open ocean, as apparent oxygen utilization has been linked to the decreases in S_275-295_ and increases in humic-like fluorescence (Catalá et al., 2015b, 2018; Martínez-Pérez et al., 2017).

### 4.4 Implications for carbon sequestration

The fertilization with nutrient-rich deep water caused an accumulation of DOC with no visible decrease during the duration of the experiment. The magnitude of the excess DOC was comparable to the carbon that sunk out of the mesocosms as POC (Baumann et al., 2021b). Thus, DOC represents a pool of major importance for carbon sequestration and its fate will affect the efficiency of artificial upwelling in this regard. While a certain degree of seasonality exists, DOC is known to accumulate in the oligotrophic waters of the North Atlantic subtropical gyre (Goldberg et al., 2010; Lomas et al., 2013; Romera-Castillo et al., 2016). This suggests that the prokaryotic communities present in such environments might have a reduced ability to consume at least part of the DOC (produced in situ or advected). DOC accumulated after artificial upwelling in oligotrophic regions may follow a similar fate. This persistent DOC could then potentially be subducted below the permanent pycnocline at different scales, including mesoscale processes (Boyd et al., 2019; Le Moigne, 2019). However, the duration of the experiment was relatively short (∼5.5 weeks) and the last deep water addition of the recurring mode was performed only a few days before the end. We were thus not able to do a comprehensive assessment of the DOC remineralization process in the medium and long-term. This, however, would be key for a proper assessment of the fate of the accumulated DOC.

Long-term nutrient addition experiments (weeks to >1 year) have shown that following large DOM accumulations, a significant fraction of it can be remineralised in the weeks following the diatom bloom. However, as much as ∼30% can remain for at least several months (Fry et al., 1996; Meon and Kirchman, 2001). The accumulated DOM would be extremely diverse in its molecular composition, following the consumption and transformation of organic molecules by prokaryotes (Hach et al., 2020; Lechtenfeld et al., 2015). The observed production of CDOM and humic-like FDOM (Figure 4 and 6) and their correlation with cumulative PHP (Figure 8) is indicative for ongoing microbial transformation of DOM (Catalá et al., 2015b; Nelson and Siegel, 2013). This has already been observed at the molecular level in other mesocosm fertilization experiments (Zark et al., 2017a). The complex web of ecological interactions between the diverse DOM pool and the prokaryotic community could help explain its potential long-term persistence (Dittmar et al., 2021). The decreasing concentrations of specific molecules due to consumption and further diversification would yield decreasing rates of degradation and lower abundances of prokaryotes capable of consuming them, resulting in the accumulation of the DOM pool as a whole (Dittmar et al., 2021). In the context of these interactions, favorable environmental conditions (including nutrient availability) and the specific suit of metabolic capabilities of the prokaryotes present in surface waters would be of key importance. Inorganic nutrients were quickly depleted following additions (Ortiz et al., 2022), which would contribute to hinder consumption of the accumulated DOM (Kritzberg et al., 2010; Letscher et al., 2015). In the recurring artificial upwelling mode this might be alleviated, as periodic pulses of inorganic nutrients are provided to the microbial community. The lesser CDOM accumulation and spectral slope change in the recurring mode might be related to this. However, even in the presence of nutrients, surface prokaryotic communities might not possess the required metabolic machinery to consume a fraction of DOM, leading to its accumulation (Sebastián et al., 2021). Other factors such as predation (Jürgens and Massana, 2008) and viral infection (Breitbart et al., 2018) of prokaryotes could also have influenced the bulk degradation rates, as the abundance of prokaryotes (Figure S9) markedly decreased during the diatom bloom collapse and the DOM accumulation period that followed, subsequently presenting another two major oscillations.

The proportions of DOM that are to be remineralised or persist in the environment after artificial upwelling need to be assessed in conjunction with the export of POC (Baumann et al., 2021b): in view of the magnitude of the DOC accumulation, depending on the proportion of DOC that is returned to inorganic carbon (and where this happens in the water column), a large amount of potential C sequestration would fail to materialize. A long monitoring period (multiple weeks, months) and tracing of DOC dynamics would be required to reliably assess the fate of DOC. Additionally, while mesocosms are a useful tool to study pelagic communities in close-to-natural conditions, they cannot reproduce the multi-dimensional physical dynamics of the real ocean. These would, however, need to be considered in order to address the CO_2_ sequestration potential of the enlarged DOM pool that we found in our experiment. The transport of the fixed carbon to the deep ocean before its remineralization is the requirement for carbon sequestration and thus needs to be examined in future work. Downwelling or convective mixing would need to be considered, as they transport DOC to the deep ocean (Baetge et al., 2020; Boyd et al., 2019). Moreover, horizontal advection would potentially limit the accumulation of DOC, thereby reducing its concentrations and making potential downwelling less effective. All these factors would need to be considered when applying artificial upwelling in natural, open-sea conditions.

## 5 Conclusions

The artificial upwelling of deep, nutrient-rich waters into the oligotrophic subtropical North Atlantic yielded marked increases in the DOM concentration and its carbon content, and shifts in DOM characteristics. The magnitude of the observed changes was mostly related to the upwelling intensity, as mesocosms subject to more intense upwelling presented higher concentrations of DOC. Increases over 70 μM for extreme treatments show the potential of artificial upwelling to transfer inorganic carbon to the dissolved organic fraction. The resulting DOC pool was as large as the POC that sunk in the mesocosms, highlighting the importance that DOC has for carbon sequestration. Upwelling intensity was also related to carbon enrichment of DOM, increases in concentration and average molecular weight of CDOM, and intensification of signals of humic-like FDOM. The generation of CDOM, and specifically humic-like FDOM, has been associated with the reworking of organic matter, suggesting ongoing transformation and molecular diversification of DOM during the experiment. The artificial upwelling method yielded partially different outcomes: while it did result in differences in DOC concentrations during the time period of the experiment, the treatments reproducing a singular upwelling event presented higher CDOM quantities and average molecular weight than recurring treatments, as well as differences in the spectral slope ratios. These differences in the CDOM pool might indicate that the singular upwelling event yielded conditions where the by-products of the microbial transformation of DOM accumulated to a greater extent than in recurring upwelling. Nonetheless, in both upwelling modes no decreases in DOM quantity were observed. This persistence could be associated with a combination of the molecular diversification of DOM due to microbial reworking, unfavorable environmental conditions (nutrient limitation) and inadequate metabolic capabilities of the prokaryotic communities in the mesocosms. While the temporal scale of the experiment leaves an open question regarding the mid-and long-term fate of the accumulated DOM (long-term persistence vs gradual remineralization), our results highlight the importance of considering DOC along POC when assessing the carbon sequestration potential of artificial upwelling. A monitoring period of multiple weeks/months would be required to reliably estimate the extent of DOM remineralization and studies in open-sea conditions would be necessary to include the effects of the physical processes involved in carbon export.

## 7 Author Contributions

Experimental concept and design: UR and JA. Execution of experiment and sample analyses: all authors. Data analysis: MGL with contributions from all authors. MGL wrote the original draft with input from all authors.

## 9 Acknowledgments

The authors thank the Plataforma Oceánica de Canarias (PLOCAN) for their extensive support throughout the experiment, including the use of their facilities and their assistance during the sampling of the mesocosms. The authors thank the captain and crew of RV James Cook for the deployment of the mesocosms and the deep water collection, and the captain and crew of the vessel J. SOCAS for helping with the second deep water collection and the recovery of the mesocosms at the end of the experiment. The authors especially thank the whole KOSMOS team (GEOMAR) for their relentless effort organizing and carrying out all the logistic and technical work necessary for the experiment, and to the biological oceanography group of the University of Las Palmas de Gran Canaria (GOB-ULPGC) for providing lab facilities and technical support. Particular thanks go to Acorayda González for her contribution to the measurement of dissolved organic carbon.

## 10 Data Availability Statement

The dataset presented in this study is available online in the PANGAEA repository (www.pangaea.de) under accession number XXXXXXXXX.

