## Supplementary material for "Artificial upwelling leads to a large increase in surface dissolved organic matter concentrations"

### 1 Supplementary Figures and Tables

#### 1.1 Supplementary Figures

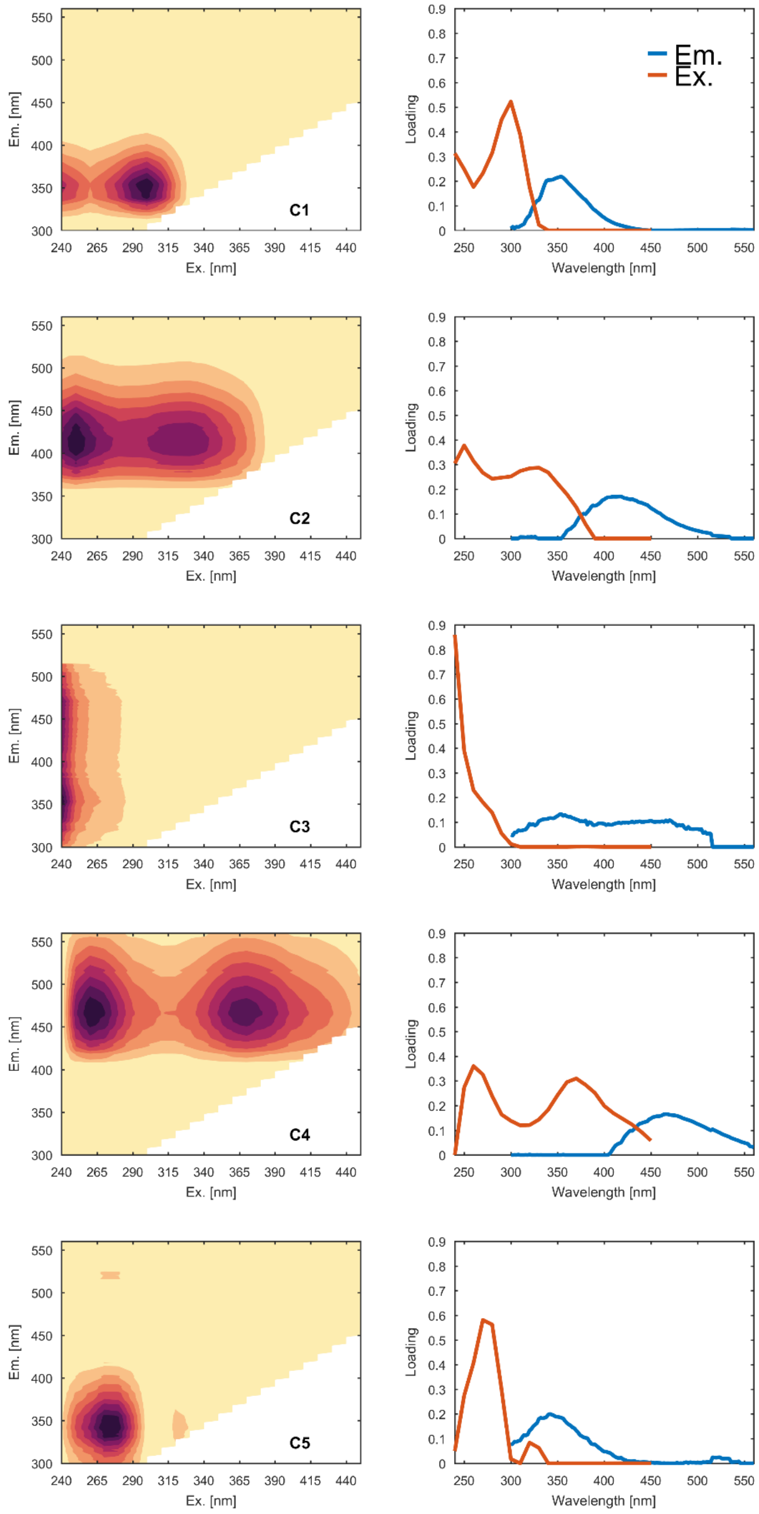

**Supplementary figure 1**. FDOM components derived from the PARAFAC analysis (C1, C2, C3, C4, C5; see Methods for details). In the left column, excitation-emission matrices of each component; in the right column, excitation (red) and emission (blue) spectra. The processed EEMs were analysed using the DOMFluor toolbox (v. 1.7; Stedmon and Bro, 2008) toolbox for Matlab (R2017a).

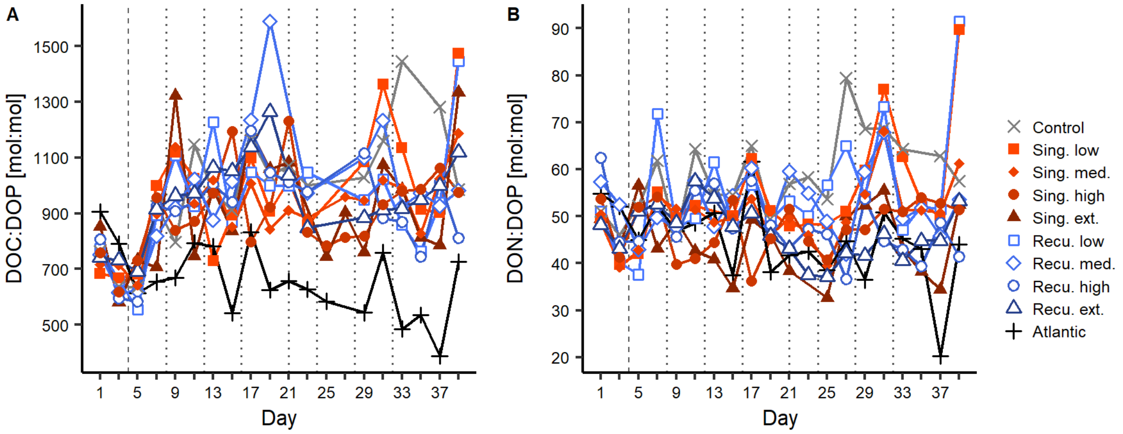


**Supplementary Figure 2**. Changes in dissolved organic matter concentrations and ratios during the experiment. Temporal evolution of A) DOC:DOP, and B) DON:DOP. Vertical lines indicate deep water additions of singular (dashed, day 4) and recurring (dashed and dotted) treatments.

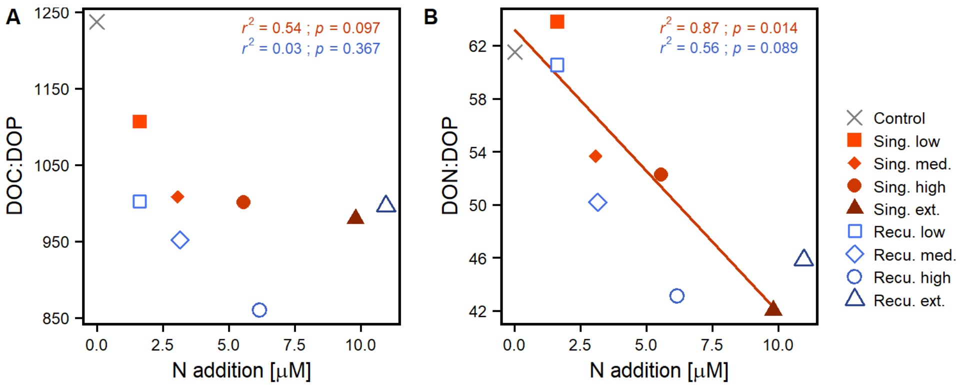


**Supplementary figure 3**. Effect of upwelling intensity on dissolved organic matter. Shown are linear regressions of A) DOC:DOP and B) DON:DOP values against upwelling intensity (N addition) per upwelling mode. DOM values were averaged after the last deep water addition (≥ day 33). The coefficient of determination ($r^2$) and p-value (p) of the regressions are included. Only the lines for significant regressions ($p < 0.05$) are displayed (see Table S3 for detailed test statistics).

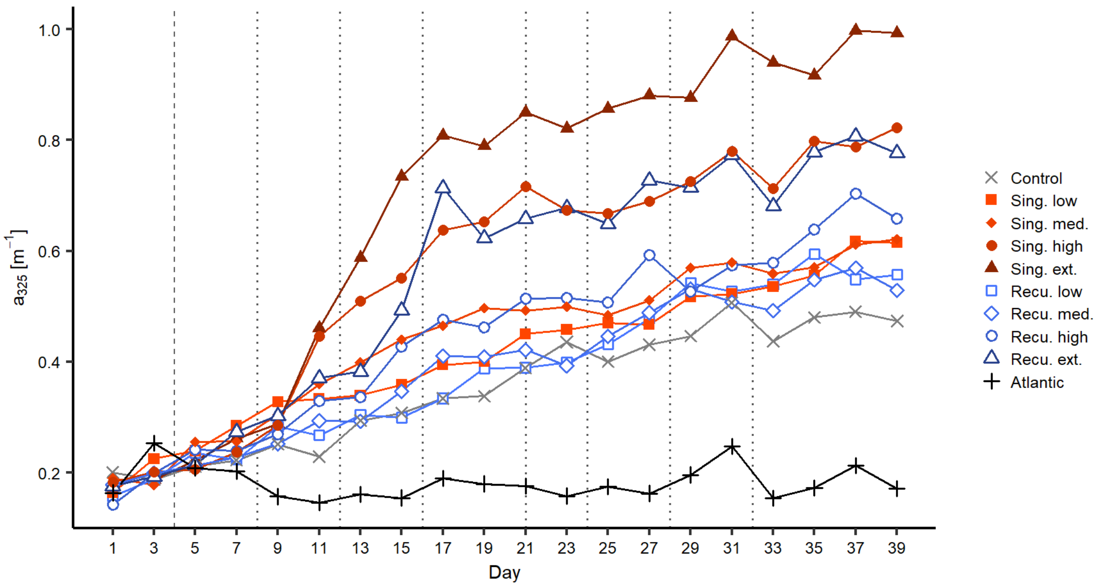


**Supplementary figure 4**. Temporal evolution of the absorption coefficient at 325 nm ($a_{325}$) during the experiment.

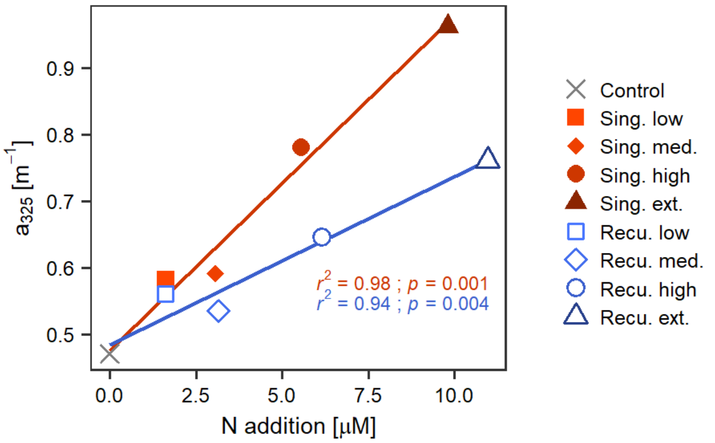


**Supplementary figure 5**. Linear regression of average values after the last deep water addition (≥ day 33) of $a_{325}$ against upwelling intensity (as N addition), per upwelling mode. The coefficient of determination ($r^2$) and p-value ($p$) of the regressions are included.

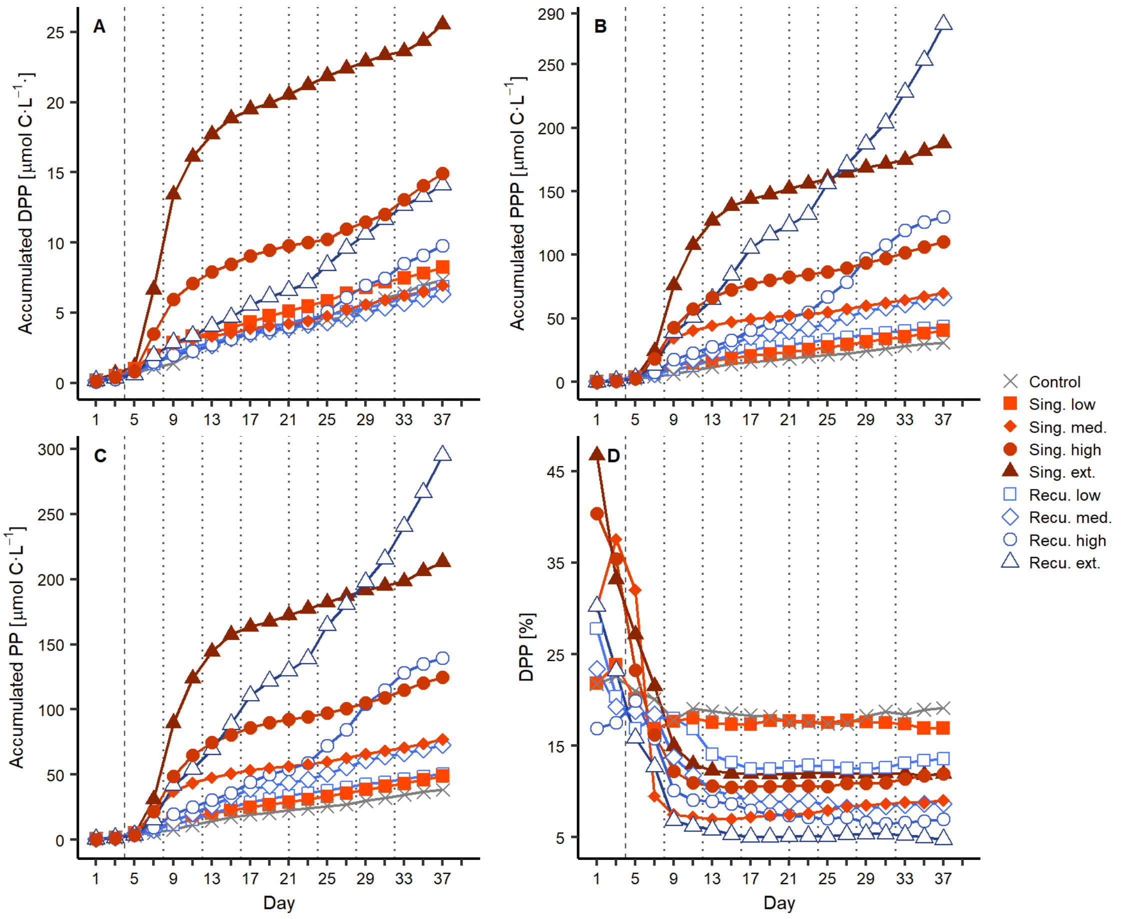


**Supplementary figure 6**. Temporal evolution of accumulated PP in the different fractions over the experiment. A) dissolved primary production (DPP), B) particulate primary production (PPP), C) total primary production (PP), and D) DPP as % of total PP.

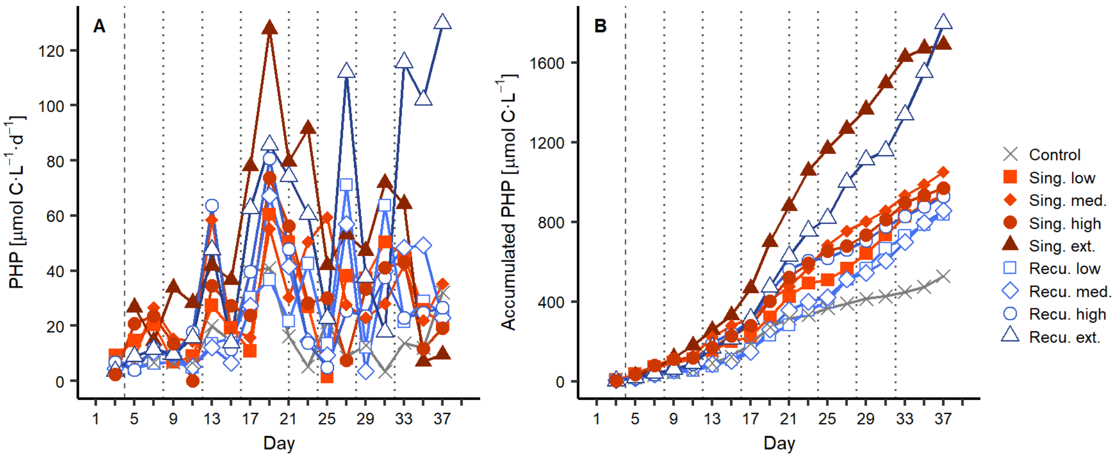

**Supplementary figure 7**. Temporal evolution of A) prokaryotic heterotrophic production (PHP) and B) its accumulated values.

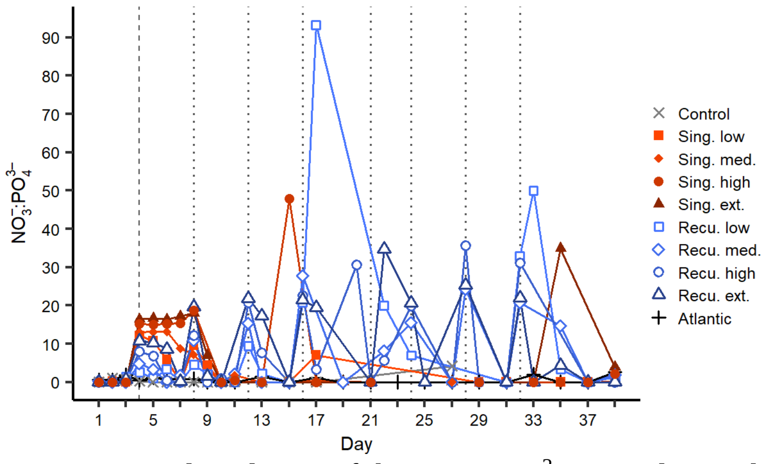


**Supplementary figure 8**. Temporal evolution of the $NO_3^-:PO_4^{3-}$ ratios during the experiment.

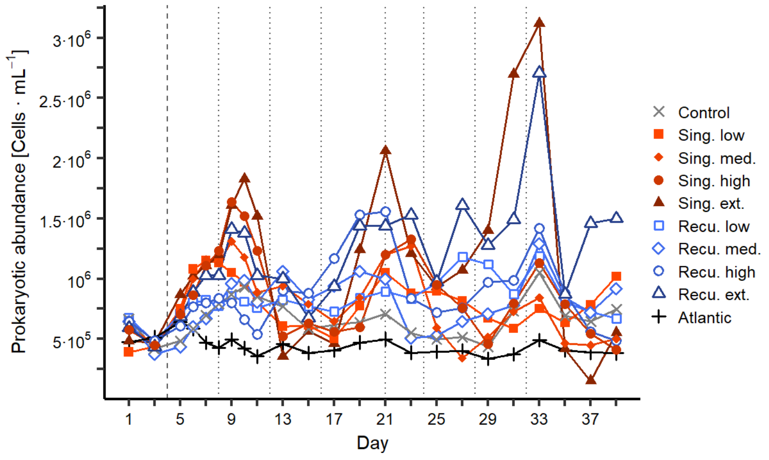


**Supplementary figure 9**. Temporal evolution of the abundance of prokaryotes during the experiment as determined by flow cytometry.

#### 1.2 Supplementary Tables

**Supplementary table 1**. Dissolved organic matter characteristics of the Deep Water employed during the artificial upwelling experiment. See Materials and methods section in the main text for details on parameters. Data from days 3-25 correspond to the first Deep Water bag (330 m) and from days 27-31 to the second Deep Water bag (280 m).

| Day | DOC [µM] | DON [µM] | DOP [µM] | DOC:DON | DOC:DOP | DON:DOP | $a_{254}$ [$m^{-1}$] | $a_{325}$ [$m^{-1}$] | $S_{275-295}$ [$\mu m^{-1}$] | $S_{350-400}$ [$\mu m^{-1}$] | $S_R$ | C1 [RU] | C2 [RU] | C4 [RU] | C5 [RU] |
|---|---|---|---|---|---|---|---|---|---|---|---|---|---|---|---|
| 3 | - | 4.36 | 0.052 | - | - | 84.5 | 1.29 | 0.278 | 24.63 | 11.33 | 2.17 | - | - | - | - |
| 7 | 82 | 3.27 | 0.04 | 25.1 | 2051 | 81.6 | 1.24 | 0.232 | 27.46 | 12.29 | 2.23 | 0.008 | 0.012 | 0.008 | 0.004 |
| 9 | 104.6 | 3.72 | 0.057 | 28.1 | 1845 | 65.6 | 1.33 | 0.233 | 27.82 | 12.59 | 2.21 | - | - | - | - |
| 11 | 116.7 | 4.37 | 0.078 | 26.7 | 1505 | 56.4 | 1.48 | 0.263 | 27.47 | 13.15 | 2.09 | 0.02 | 0.015 | 0.01 | 0.008 |
| 15 | 184.6 | 6.87 | 0.129 | 26.9 | 1432 | 53.3 | 1.9 | 0.44 | 23.39 | 12.6 | 1.86 | 0.035 | 0.016 | 0.013 | 0.011 |
| 17 | - | 6.84 | 0.149 | - | - | 45.8 | - | - | - | - | - | - | - | - | - |
| 19 | 178.8 | 4.94 | 0.119 | 36.2 | 1500 | 41.5 | 2.4 | 0.573 | 23.92 | 13 | 1.84 | 0.017 | 0.019 | 0.015 | 0.016 |
| 25 | - | 3.42 | 0.045 | - | - | 75.9 | 2.2 | 0.507 | 23.83 | 12.8 | 1.86 | 0.032 | 0.024 | 0.017 | 0.018 |
| 27 | 53.3 | 4.54 | 0.053 | 11.7 | 996 | 85 | 1.05 | 0.195 | 27.24 | 11.09 | 2.46 | - | - | - | - |
| 29 | - | 3.81 | 0.012 | - | - | 322.3 | - | - | - | - | - | - | - | - | - |
| 31 | 71.4 | 4.68 | 0.052 | 15.3 | 1371 | 89.8 | 1.08 | 0.191 | 28.19 | 12.33 | 2.29 | 0.011 | 0.01 | 0.007 | 0.003 |

**Supplementary table 2**. Characteristics of the FDOM components derived from the PARAFAC analysis, along with analogous fluorophores previously described in the literature. Components are named according to the wavelength of their emission maximum. FDOM fluorophores from the literature were identified making use of the OpenFluor database (Murphy et al., 2014), applying threshold values of the Tucker Congruence Coefficients (TCC) of 0.95 for both excitation and emission spectra. Wavelengths between parentheses represent secondary maxima.

| ***This study*** | | | ***Similar fluorophores in literature*** | | | | | |
|---|---|---|---|---|---|---|---|---|
| Comp. | Excitation max [nm] | Emission max [nm] | Study | Comp. name | Excitation max [nm] | Emission max [nm] | $TCC_{ex\cdot em}$ | Description |
| C1 | 300 (240) | 354 | Chen et al. (2018) | $C_{305/344}$ | 305 | 344 | 0.9829 | Amino acid-like. Peak T (Coble, 1996). |
| | | | Catalá et al. (2015) | C3 | 290 | 340 | 0.9827 | |
| | | | Amaral et al. (2020) | C2 | 300 | 359 | 0.9670 | |
| | | | Amaral et al. (2016) | C3 | 300 (<250) | 340 | 0.9930 | |
| C2 | 250 (330) | 410 | Yamashita et al. (2011b) | C1 | <250 (320) | 422 | 0.9853 | Humic-like, peak M (Coble, 1996). Positively correlated with AOU |
| | | | Catalá et al. (2015) | C2 | 320 | 400 | 0.9774 | |
| | | | Yamashita et al. (2011a) | C1 | <260 (320) | 425 | 0.9774 | |
| | | | Chen et al. (2018) | $C_{<260(305)/404}$ | <260 (305) | 404 | 0.9721 | |
| | | | Amaral et al. (2020) | C1 | 270 (320) | 411 | 0.9715 | |
| C3 | <240 | 330-472 | Murphy et al. (2008) | P4 | <260 | - | 0.9248 | Low excitation maxima, broad emission spectrum. Origin unknown. Potential fluorometer artifact. |
| | | | Yamashita et al. (2010) | C3 | <260 | - | 0.9197 | |
| C4 | 260 (370) | 466 | Chen et al. (2018) | $C_{<260(365)/476}$ | <260 (365) | 476 | 0.9815 | Humic-like, mixture of A and C peaks (Coble, 1996). High aromaticity. |
| | | | Yamashita et al. (2011a) | C2 | 365 (<260) | 475 | 0.9658 | |
| | | | Yamashita et al. (2010) | C1 | <260 (370) | 466 | 0.9523 | |
| | | | Dainard et al. (2015) | C2 | <260 (370) | 475 | 0.9630 | |
| C5 | 270 | 342 | Lapierre and Del Giorgio (2014) | C6 | 275 | 334 | 0.9517 | Amino acid-like. peak T |
| | | | Chen et al. (2018) | C5 | 275 | 338 | 0.9421 | |
| | | | Osburn et al. (2015) | C7 | 280 | 340 | 0.9551 | |
| | | | Wünsch et al. (2015) | C2 | 269 | 340 | 0.9577 | Indole |

**Supplementary table 3**. Summary of results from the linear regressions of DOM variables versus upwelling intensity (as N addition, in µM), per upwelling mode. Slope and intercept values are presented along 95% confidence intervals. $r^2$ is the coefficient of determination of the regression, *p* is the p-value of the regression. Significant regressions are highlighted in *italic* ($p < 0.05$) and **bold** ($p < 0.01$).

| | **Upwelling mode** | **Slope** | **Intercept** | $r^2$ | *p* |
|---|---|---|---|---|---|
| DOC [µM] | Singular | 5.32 ± 1.98 | 89.8 ± 10.4 | 0.95 | **0.003** |
| | Recurring | 4.77 ± 1.54 | 91.91 ± 8.98 | 0.96 | **0.002** |
| DON [µM] | Singular | 0.111 ± 0.045 | 5.28 ± 0.24 | 0.94 | **0.004** |
| | Recurring | 0.121 ± 0.075 | 5.34 ± 0.44 | 0.87 | *0.014* |
| DOP [µM] | Singular | 0.0075 ± 0.0014 | 0.0789 ± 0.0076 | 0.99 | **< 0.001** |
| | Recurring | 0.0060 ± 0.0054 | 0.090 ± 0.032 | 0.74 | *0.039* |
| DOC:DON [mol:mol] | Singular | 0.51 ± 0.46 | 17.39 ± 2.41 | 0.74 | *0.039* |
| | Recurring | 0.37 ± 0.37 | 17.56 ± 2.19 | 0.7 | *0.05* |
| DOC:DOP [mol:mol] | Singular | -22.7 ± 30.2 | 1157.5 ± 159.3 | 0.54 | 0.097 |
| | Recurring | -16.8 ± 50.5 | 1083.2 ± 295.1 | 0.03 | 0.367 |
| DON:DOP [mol:mol] | Singular | -2.13 ± 1.30 | 63.21 ± 6.85 | 0.87 | *0.014* |
| | Recurring | -1.59 ± 2.05 | 59.2 ± 11.9 | 0.56 | 0.089 |
| $a_{254}$ [$m^{-1}$] | Singular | 0.095 ± 0.025 | 2.21 ± 0.13 | 0.97 | **0.001** |
| | Recurring | 0.050 ± 0.016 | 2.235 ± 0.096 | 0.96 | **0.002** |
| $a_{325}$ [$m^{-1}$] | Singular | 0.050 ± 0.013 | 0.475 ± 0.067 | 0.98 | **0.001** |
| | Recurring | 0.025 ± 0.010 | 0.483 ± 0.059 | 0.94 | **0.004** |
| $S_{275-295}$ [$\mu m^{-1}$] | Singular | -0.68 ± 0.21 | 25.85 ± 1.11 | 0.96 | **0.002** |
| | Recurring | -0.33 ± 0.27 | 25.65 ± 1.57 | 0.78 | *0.029* |
| $S_{350-400}$ [$\mu m^{-1}$] | Singular | 0.084 ± 0.145 | 15.10 ± 0.77 | 0.37 | 0.163 |
| | Recurring | -0.14 ± 0.05 | 15.13 ± 0.30 | 0.95 | **0.003** |
| $S_R$ | Singular | -0.052 ± 0.020 | 1.71 ± 0.11 | 0.94 | **0.004** |
| | Recurring | -0.0072 ± 0.0182 | 1.70 ± 0.11 | 0.12 | 0.299 |
| $C_{354}$ [RU] | Singular | 0.0045 ± 0.0072 | 0.027 ± 0.038 | 0.42 | 0.143 |
| | Recurring | -0.00034 ± 0.00173 | 0.040 ± 0.010 | -0.18 | 0.578 |
| $C_{410}$ [RU] | Singular | 0.00093 ± 0.00017 | 0.01170 ± 0.00090 | 0.99 | **< 0.001** |
| | Recurring | 0.00064 ± 0.00029 | 0.0121 ± 0.0017 | 0.92 | **0.006** |
| $C_{466}$ [RU] | Singular | 0.00033 ± 0.00028 | 0.0069 ± 0.0015 | 0.77 | *0.033* |
| | Recurring | 0.00025 ± 0.00037 | 0.0071 ± 0.0022 | 0.47 | 0.124 |
| $C_{342}$ [RU] | Singular | 0.00050 ± 0.00032 | 0.0106 ± 0.0017 | 0.85 | *0.016* |
| | Recurring | 0.00080 ± 0.00075 | 0.0121 ± 0.0044 | 0.73 | *0.042* |
